# The intestinal microbiome and *Cetobacterium somerae* inhibit viral infection through TLR2-type I IFN signaling axis in zebrafish

**DOI:** 10.1101/2024.05.25.595869

**Authors:** Hui Liang, Ming Li, Jie Chen, Wenhao Zhou, Dongmei Xia, Qianwen Ding, Yalin Yang, Zhen Zhang, Chao Ran, Zhigang Zhou

## Abstract

Accumulated evidences demonstrate that intestinal microbiome can inhibit viral infection. However, our knowledge of the signaling pathways and specific commensal microbes that mediate the antiviral response is limited. Here, a rhabdoviral infection model in zebrafish allows us to investigate the modes of action of microbiome-mediated antiviral effect. We observed that antibiotics-treated and germ-free zebrafish exhibited greater viral infection. Mechanistically, depletion of the intestinal microbiome alters TLR2-Myd88 signaling, and blunts neutrophil response and type I interferon (IFN) production. Moreover, a single commensal bacterium, *Cetobacterium somerae*, recapitulated TLR2– and type I IFN-dependent antiviral effect of the microbiome in gnotobiotic zebrafish, and *C. somerae* exopolysaccharides (CsEPS) was the effector molecule that engaged TLR2 to mediate antiviral function. Together, our results suggest a conserved role of intestinal microbiome in regulating type I IFN response among vertebrates, and reveal that the intestinal microbiome inhibits viral infection through a CsEPS-TLR2-type I IFN signaling axis in zebrafish.

## INTRODUCTION

Vertebrates harbor a large number of commensal bacteria in the intestine. The intestinal microbiota play an important role in the health and disease of host, both locally and systemically (*1*, *2*). During the past decade, accumulated evidences shed light on the regulatory role of intestinal microbiota in viral infection (*3–5*). The microbiota can either stimulate or inhibit viral infection, depending on the viruses investigated (*5*, *6*). Notably, intestinal microbiota has been shown to limit mammalian infection of different viruses (*7-12*). The underlying mechanism was associated with commensal bacteria-mediated priming of type I interferon (IFN) response in a number of studies (*8–11*). Till now, studies about the influence of intestinal microbiota on viral infection are mainly conducted in mammalian model. Recent work demonstrated that intestinal microbiota improved resistance of chicken against nephropathogenic infectious bronchitis virus (*12*), suggesting a conserved role of microbiota in regulating viral infection. However, the influence of commensal bacteria on viral infection in lower vertebrates is largely unknown. Furthermore, despite the advances in this field, our knowledge about the identity of specific commensal bacteria that primes the antiviral immunity and the underlying molecular mechanism is still limited (*2*, *13*).

Zebrafish have emerged as a powerful animal model for study of vertebrate-microbiota interactions (*14*). Gnotobiotic zebrafish revealed evolutionarily conserved responses to the gut microbiota compared with mice (*15*). Studies using gnotobiotic zebrafish have given rise to important insights into the assembly and function of vertebrate microbiome (*14*, *16–20*). Zebrafish possess innate and adaptive immune system similar to that of mammals, and rely solely on innate immunity at least within 3 weeks of its life (*21*). Similar with mammals, virus-induced IFNs are key components of antiviral immunity in zebrafish. The type I IFNs in zebrafish comprise IFNΦ1-4, which are divided into two groups: I (IFNΦ1 and Φ4) and II (IFNΦ2 and Φ3), and are recognized by different heterodimeric receptors: CRFB1/CRFB5 and CRFB2/CRFB5, respectively (*22–25*). Spring viremia of carp virus (SVCV) is an aquatic single stranded RNA virus that belongs to *Rhabdoviridae* family (*26*). Zebrafish are susceptible to SVCV, and zebrafish-SVCV infection model has been widely used to study the regulation of antiviral innate immunity in vertebrate (*27–29*).

In this study, we investigated the impact of intestinal microbiome on SVCV infection in zebrafish. We observed that oral antibiotics-treated and germ-free zebrafish exhibited greater viral infection by SVCV, supporting a conserved inhibitory effect of microbiome on viral infection in this lower vertebrate model. These phenotypes were attributable to microbiome-mediated priming of type I IFNs response, and neutrophils were the immune cells mediating the antiviral function of microbiome. The microbiota-mediated effect relied on toll-like receptor 2 (TLR2)-Myd88 signaling pathway. Moreover, we identified a specific commensal bacterium dictating the antiviral effect of the microbiome. Indeed, colonization of GF zebrafish with the commensal bacterial species, *Cetobacterium somerae* (*C. somerae*), inhibited SVCV infection in a mode involving TLR2-type I IFN signaling and neutrophils. We also determined that the exopolysaccharides of *C. somerae* are the effector component that engages TLR2 to inhibit viral infection.

## RESULTS

### Depletion of the intestinal microbiota modulates the resistance of zebrafish against SVCV infection

We feed adult zebrafish control feed or feed supplemented with antibiotics cocktail (ABX) for 2 weeks to deplete the microbiota. Then zebrafish were challenged by SVCV through intraperitoneal injection. The mortality rate was observed and documented for 10 days post infection. The mortality of ABX group was significantly higher than control (Fig. 1A). Furthermore, viral replication was measured by *q*PCR analysis of the transcript of SVCV N protein in infected animal tissues. Consistent with the mortality rate, viral replication in liver, spleen, and kidney of zebrafish at 1 and 2 days post infection was higher in ABX-treated fish compared with control fish (Fig. 1B). Together, these results indicate that the intestinal microbiota regulates antiviral resistance of zebrafish.

**Fig. 1.**
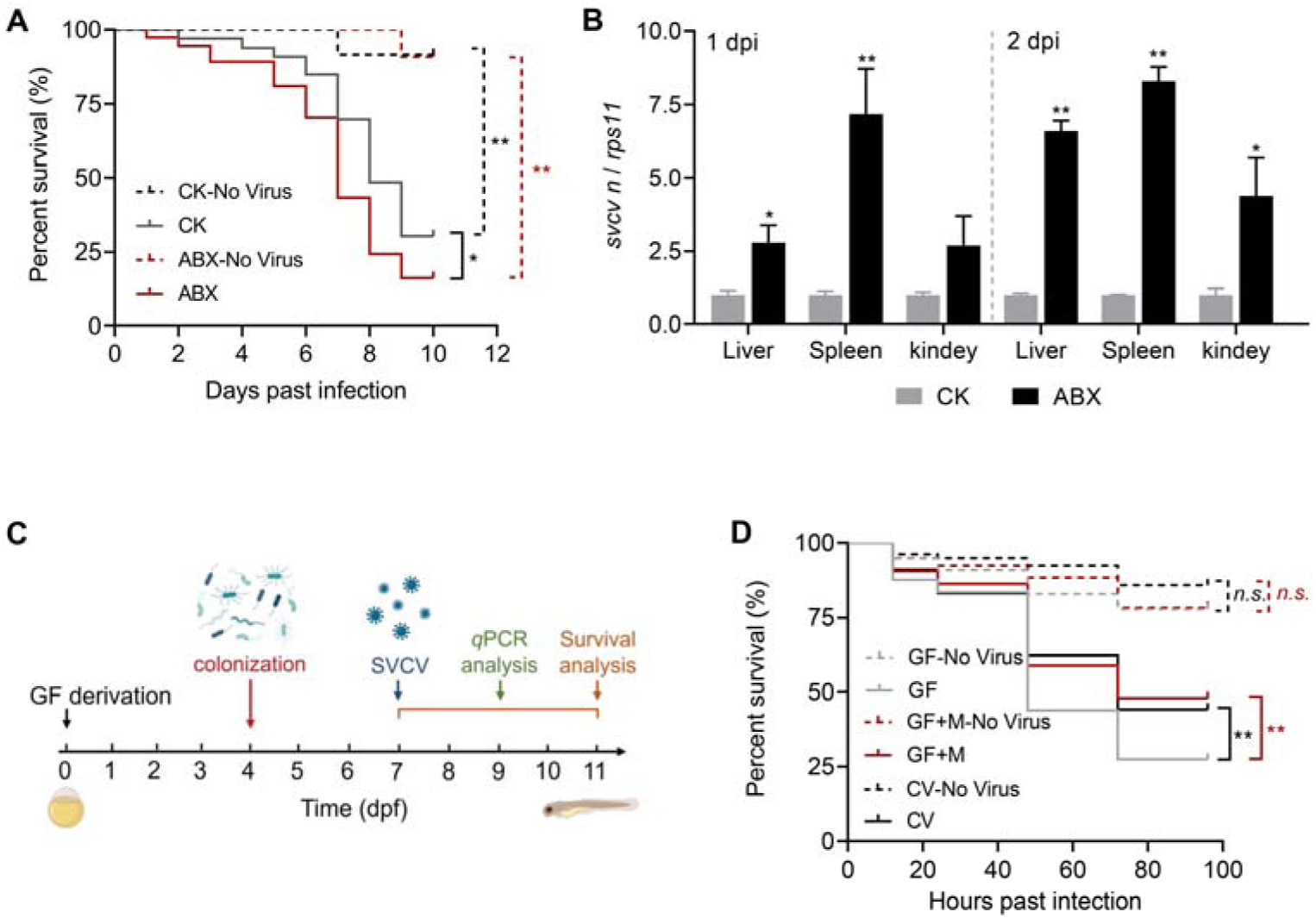
Depletion of intestinal microbiome enhances SVCV infection in zebrafish. (**A**) Survival curve of mock or SVCV-infected adult zebrafish fed control or ABX diet (No Virus groups: n=11-12; SVCV groups: n=39-40). (**B**) Viral replication in the liver, spleen, and kidney of SVCV-infected adult zebrafish fed control or ABX diet (n=4, pool of 6 fish per sample). (**C**) Schematic representation for gnotobiotic zebrafish experiment. (**D**) Survival curve of mock or SVCV-infected GF, conventional, and conventionalized zebrafish (n = 80). (A and D) log-rank test; (B) unpaired *t* test. \**p*< 0.05, \*\**p*< 0.01, *n.s.*, not significant.

To confirm a role for intestinal microbiota in modulating SVCV infection, we further conducted experiments in gnotobiotic zebrafish model. Germ-free (GF) zebrafish, conventional zebrafish, and GF zebrafish colonized by microbiota from conventionally raised adult zebrafish at 4 dpf (conventionalized) were challenged by SVCV at 7 dpf, and the mortality was monitored (Fig. 1C). The results showed that the mortality of germ free group was higher than both conventional or conventionalized counterparts (Fig. 1D), which confirms the antiviral effect of microbiota in zebrafish.

### Intestinal microbiota depletion impairs the type I IFN response to SVCV infection

To investigate the potential mechanism of the antiviral effect of microbiota, we firstly evaluated the type I IFN response in germ-free versus conventionalized zebrafish. The results showed that the expression of IFNΦ1 and IFNΦ3 was higher in conventionalized zebrafish compared with germ free counterparts (Fig. 2A). We also investigated the type I IFN response in adult zebrafish fed control diet or diet supplemented with antibiotics. Consistently with the results in gnotobiotic fish, adult zebrafish with depleted microbiota exhibited impaired expression of IFNΦ1 and IFNΦ3 in liver, spleen, and kidney at 1 and 2 days post SVCV infection (Fig. 2, B and C). To eliminate potential confounding effect of viral replication on IFN response, we treated adult zebrafish with poly(I:C) by intraperitoneal injection. We found similar impairment of IFNΦ1 and IFNΦ3 expression in antibiotics fed zebrafish compared with control at 1 and 2 days post poly(I:C) inoculation (fig. S1). The concordance of finding in different experimental models indicated that the microbiota plays an important role in stimulation of type I IFN response in zebrafish post viral infection.

**Fig. 2.**
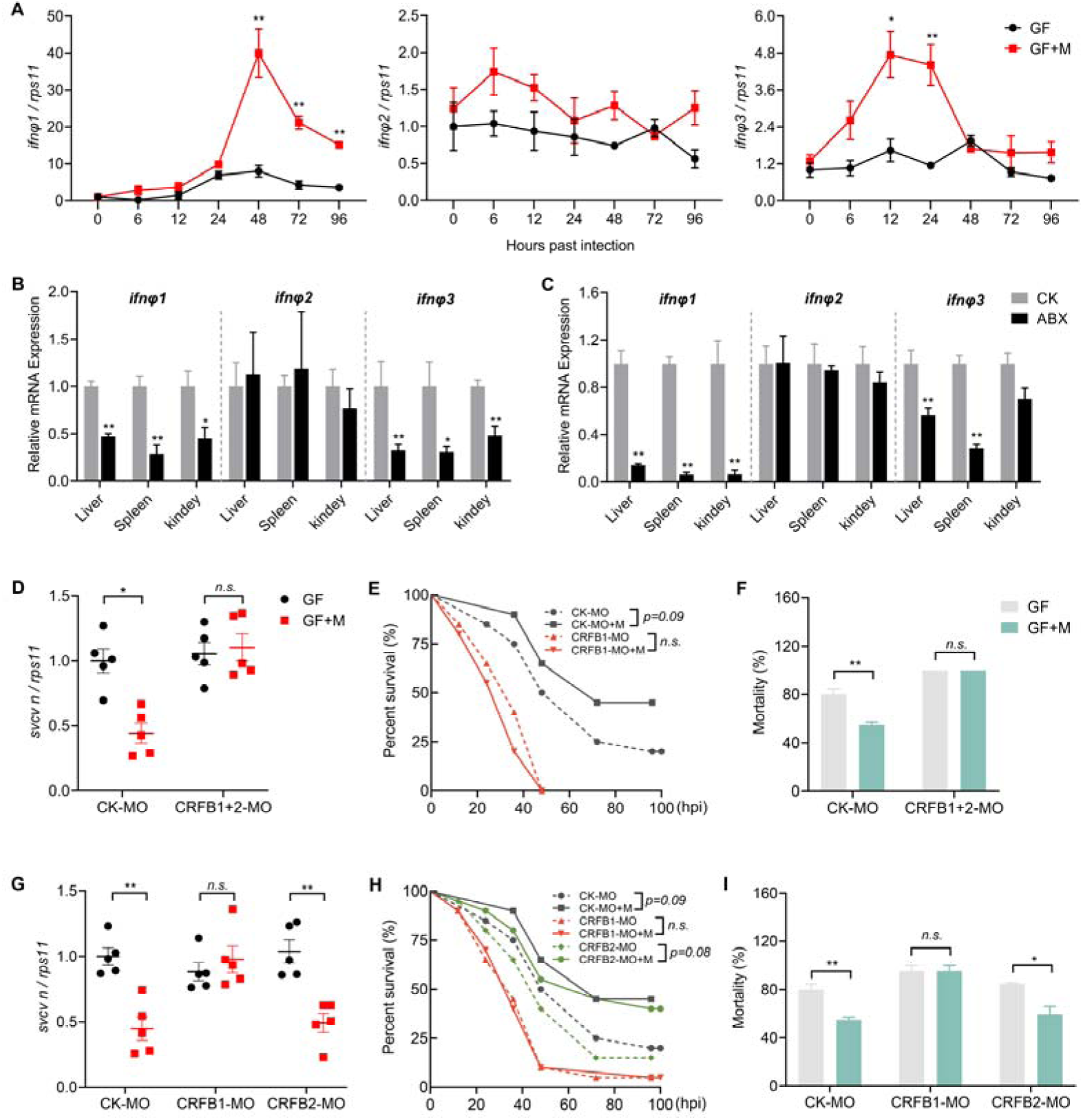
Intestinal microbiome depletion impairs type I IFN response to SVCV infection. (**A**) Expression of IFNΦ1, IFNΦ2, and IFNΦ3 in SVCV-infected GF or conventionalized zebrafish at different time points (n = 3, pool of 30 zebrafish larvae per sample). (**B**-**C**) Expression of IFNΦ1, IFNΦ2, and IFNΦ3 in the liver, spleen, and kidney of adult zebrafish after 1 (B) and 2 (C) days post SVCV infection (n = 4, pool of 6 fish per sample). (**D**-**F**) Effect of morpholino-mediated knockdown of type I IFN receptors on SVCV infection. GF and conventionalized zebrafish were treated with control morpholino (CK-MO) or a mixture of CRFB1 and CRFB2 morpholino (CRFB1+2 MO) and subjected to SVCV infection. (D) Viral replication at 48 hpi. (n = 5, pool of 30 zebrafish larvae per sample), (E) Survival curve (n=20), (F) Mortality at 96 hpi (n=3). (**G**-**I**) Effect of morpholino-mediated knockdown of group I or II type I IFN signaling on SVCV infection. GF and conventionalized zebrafish were treated with control morpholino (CK-MO), CRFB1 morpholino (CRFB1-MO), or CRFB2 morpholino (CRFB2-MO) and subjected to SVCV infection. (G) Viral replication at 48 hpi (n = 5, pool of 30 zebrafish larvae per sample), (H) Survival curve (n=20), (I) Mortality at 96 hpi (n=3). (A-D, F, G, I) unpaired *t* test; (E and H) log-rank test. \**p*< 0.05, \*\**p*< 0.01, *n.s.*, not significant.

To investigate whether the protective effect of microbiota against viral infection is mediated through type I IFN signaling, we knocked down receptors for all type I IFNs with vivo-morpholino directed to CRFB1 and CRFB2, and infected GF and conventionalized control or CRFB1+2 morphant fish with SVCV. Viral replication in gnotobiotic zebrafish was evaluated by *q*PCR analysis of SVCV N protein mRNA at 48 hpi (Fig. 1C). The results showed that CRFB1+2 knockdown abrogated the antiviral effect of microbiota as manifested by both viral replication and mortality (Fig. 2, D to F, fig. S2, A and B), indicating that the antiviral effect of intestinal microbiota depends on type I IFN response in zebrafish. In addition to type I IFN, other types of IFNs were also reported to contribute to the protection effect of microbiota against viral infection in some studies (*7*, *13*). To investigate the potential involvement of type II and IV IFNs (*30*) in the microbiota-mediated effect, we detected the expression of IFNγ, IFNγrel, IFNυ in GF and conventionalized zebrafish post SVCV infection. We found that there was no difference in the expression of type II and IV IFNs between GF zebrafish and the microbiota-colonized counterparts at 24 and 48 hpi (fig. S3), suggesting that the microbiota-mediated protective effect on viral infection does not involve type II and IV IFN response. We also separately knocked down CRFB1 and CRFB2 to evaluate the relative contribution of group I and II type I IFNs to the antiviral effect of microbiota. The results showed that knocking down CRFB1 blocked the effect of microbiota, while CRFB2 knockdown made no difference (Fig. 2, G to I, fig. S2, C and D). This suggests that the antiviral effect of microbiota is mainly dependent on group I IFN. Group 1 IFN includes IFNΦ1 and IFNΦ4, but IFNΦ4 had little activity (*31*), indicating that IFNΦ1 is the key interferon that mediates the antiviral effect of microbiota in zebrafish. Therefore, we mainly detected the expression of IFNΦ1 in the following experiments.

### Microbiota-mediated antiviral effect depends on neutrophil response

We then studied the cellular immunity involved in the antiviral effect of microbiota. At the larval stage of zebrafish, cellular immunity only consists of myeloid cells, and neutrophils and macrophages are the main effector cells. Therefore, we firstly assessed neutrophil recruitment and activation in GF and conventionalized *Tg* (*mpx:EGFP*) zebrafish post SVCV infection. The results showed that there was a marginal increase of neutrophil numbers in GF fish after viral infection. In contrast, the number was significantly increased in conventionalized counterparts, indicating that the intestinal microbiota enhances the neutrophil response post viral infection (Fig. 3, A and B). Similarly, we conducted experiments in *Tg* (*mpeg1:EGFP*) zebrafish to assess macrophage activation. The results showed that both GF and conventionalized zebrafish exhibited a similar increase of the macrophage numbers post viral infection (Fig. 3, C and D), indicating that the intestinal microbiota is dispensable for macrophage activation in response to viral infection.

**Fig. 3.**
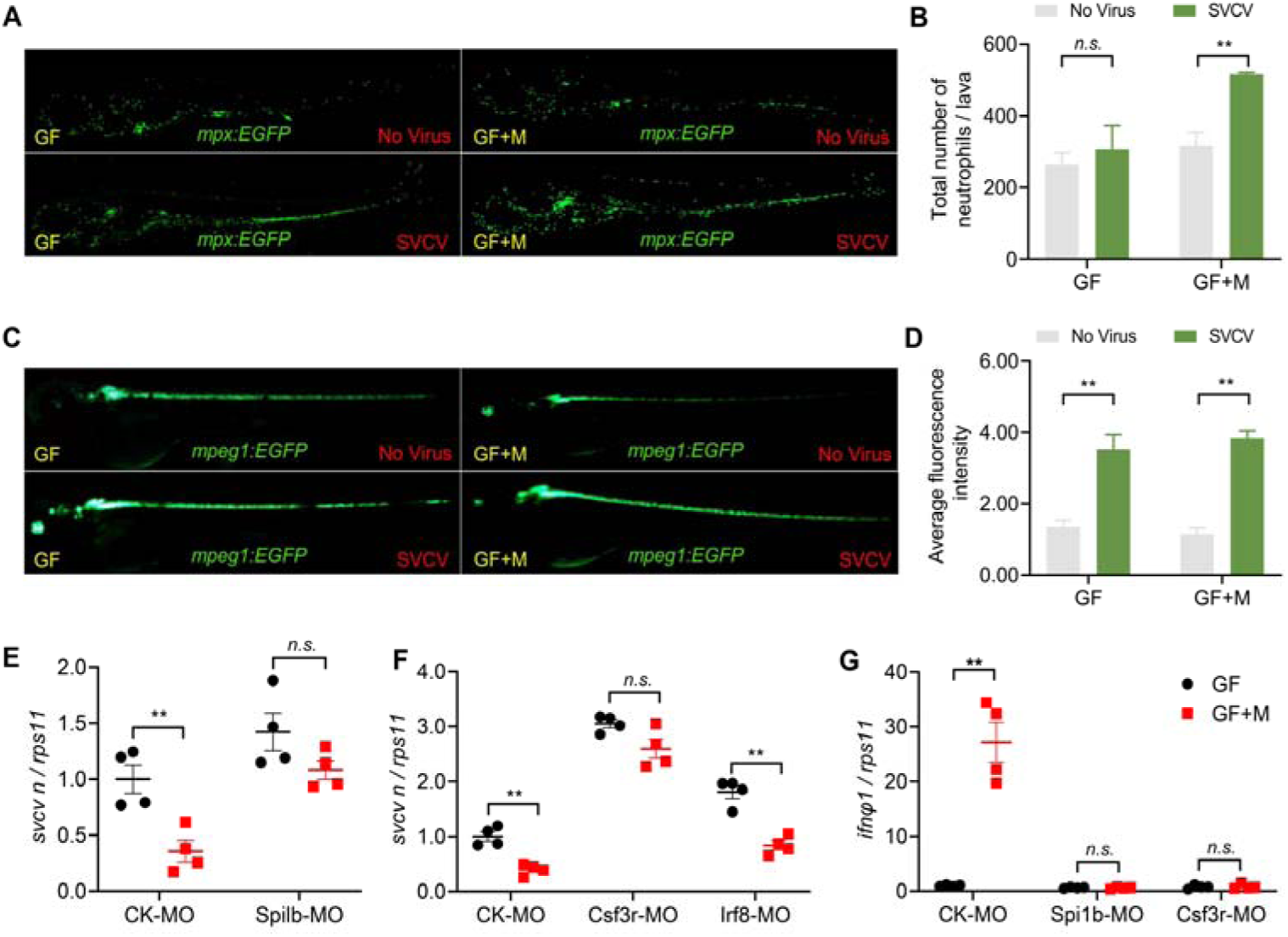
Neutrophil responses to SVCV infection are impaired in the absence of intestinal microbiome. (**A**-**D**) Neutrophils (A) and macrophages (C) were imaged in mock or SVCV-infected GF or conventionalized transgenic zebrafish at 48 hpi. Scale bar, 500 μm. The number of neutrophils (B) and macrophages (D) were analyzed (n=3). (**E**-**F**) Effect of myeloid cell depletion (Spi1b MO) (E) or selective depletion of neutrophils (Csf3r MO) or macrophages (Irf8 MO) (F) on viral replication in GF or conventionalized zebrafish at 48 hpi. (n = 4, pool of 30 zebrafish larvae per sample). (**G**) Effect of myeloid cell depletion (Spi1b MO) or selective depletion of neutrophils (Csf3r MO) on IFNΦ1 expression in GF or conventionalized zebrafish at 48 hpi (n = 4, pool of 30 zebrafish larvae per sample). (B, D, E-G) unpaired *t* test. \**p*< 0.05, \*\**p*< 0.01, *n.s.*, not significant.

Furthermore, we investigated the role of immune cells in the microbiota-mediated antiviral effect. Firstly, we blocked myelopoiesis by knocking down Spi1b, resulting in reduction of both neutrophil and macrophage populations. Spi1b knockdown blocked the antiviral effect of microbiota (Fig. 3E), indicating that the protective effect of microbiota against viral infection depends on the function of myeloid cells. Knockdown or mutation of Csf3r affected neutrophil populations in zebrafish larvae (*31*, *32*), and Irf8 is critical for macrophage development in zebrafish (*33*). To distinguish the roles of the two leukocyte types, we attempted to deplete neutrophils and macrophages by knocking down Csf3r and Irf8, respectively (fig. S4). We found that Csf3r knockdown blocked the protective effect of microbiota against viral infection, while Irf8 knockdown showed no effect (Fig. 3F). We hypothesized that neutrophils played a role in IFN production. Indeed, Spi1b and Csf3r knockdown abrogated the effect of microbiota to stimulate IFNΦ1 expression post viral infection (Fig. 3G), indicating that the stimulatory effect of microbiome on type I IFN response was mediated by neutrophils. Together, these data are consistent with the results of neutrophil/macrophage activation in conventionalized versus GF zebrafish after viral infection, supporting that the intestinal microbiota-mediated restrictive effect on viral infection depends on neutrophils, but not macrophages.

### The intestinal microbiota signals through TLR2-Myd88 to limit viral infection

We further studied whether specific PRR(s) mediated the antiviral effect of microbiota. We found that Myd88 knockdown abrogated the protective effect of microbiota against viral infection (Fig. 4A, fig. S5). Myd88 is the adaptor protein of multiple TLRs (*34*). We hypothesized that specific TLR(s) recognized the microbiota-derived signals and triggered the antiviral response to limit SVCV infection. We used vivo morpholinos to knock down one of five TLRs (TLR2, TLR3, TLR4ba, TLR7, TLR9), and infected GF and conventionalized control or TLR-morphant zebrafish with SVCV. The results showed that knockdown of TLR3, TLR4ba, TLR7 and TLR9 did not impair the protective effect of microbiota against viral infection (Fig. 4B). In contrast, the microbiome-mediated reduction of viral infection was abrogated in TLR2-knockdowned zebrafish (Fig. 4, C to E), suggesting a specific role of TLR2 in recognizing the microbiota-derived ligand(s) that stimulate antiviral response. Together, these data indicate that the intestinal microbiota requires TLR2-Myd88 signaling to restrict viral infection in zebrafish.

**Fig. 4.**
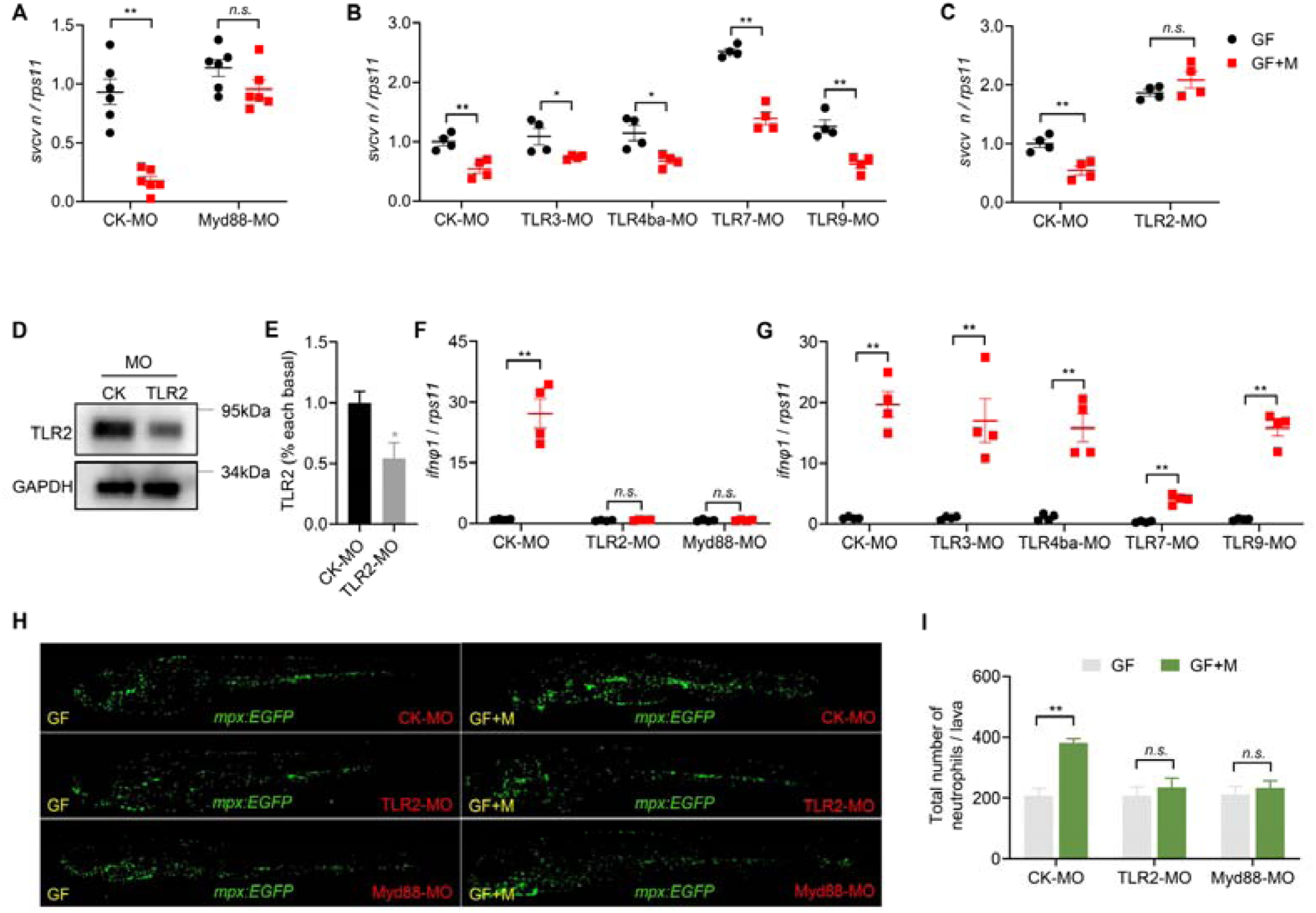
The antiviral effect of intestinal microbiota requires TLR2 and Myd88 signaling. (**A**) Effect of Myd88 knockdown on SVCV infection in GF or conventionalized zebrafish at 48 hpi (n = 6, pool of 30 zebrafish larvae per sample). (**B**) Effect of morpholino-mediated knockdown of TLR3, TLR4ba, TLR7 and TLR9 on SVCV infection in GF or conventionalized zebrafish at 48 hpi (n = 4, pool of 30 zebrafish larvae per sample). (**C**) Effect of TLR2 knockdown on SVCV infection in GF or conventionalized zebrafish at 48 hpi (n = 4, pool of 30 zebrafish larvae per sample). (**D**-**E**) TLR2 morpholino diminished TLR2 protein expression in zebrafish larvae (n=3, pool of 30 zebrafish larvae per sample). (**F**-**G**) Effect of morpholino-mediated knockdown of TLR2, Myd88 (F), TLR3, TLR4ba, TLR7 and TLR9 (G) on IFNΦ1 expression in GF or conventionalized zebrafish at 48 hpi (n = 4, pool of 30 zebrafish larvae per sample). (**H**-**I**) Effect of morpholino-mediated knockdown of TLR2 and Myd88 on neutrophil response in GF or conventionalized *Tg* (*mpx:EGFP*) zebrafish at 48 hpi. (H) Confocal imaging of SVCV-infected *Tg* (*mpx:EGFP*) zebrafish. Scale bar, 500 μm. (I) Neutrophil numbers (n=3). (A, B, C, E, F, G, I) unpaired *t* test. \**p*< 0.05, \*\**p*< 0.01, *n.s.*, not significant.

Furthermore, we evaluated the role of TLR2-Myd88 signaling in microbiota-mediated effect on type I IFN response. We found that knockdown of TLR2 and Myd88, but not any of the other TLRs, abrogated the effect of microbiota to stimulate IFNΦ1 expression post viral infection (Fig. 4, F and G). We also found that conventionalized zebrafish stimulated IFNΦ1 expression compared with GF counterparts in response to poly(I:C) treatment, and TLR2/Myd88 knockdown abrogated the effect (fig. S6), suggesting that TLR2-Myd88 signaling-mediated action of microbiome enhances canonical type I IFN signaling pathway against RNA viruses, and this mode of action is not limited to SVCV infection. To define whether TLR2-Myd88 signaling contributed to the microbiome-mediated neutrophil response, we used vivo-morpholinos to knock down TLR2 and Myd88 in *Tg* (*mpx:EGFP*) zebrafish and enumerated the neutrophil numbers in GF and conventionalized control or TLR2/Myd88 morphants after SVCV infection. Interestingly, we observed that TLR2 and Myd88 knockdown blocked microbiota-mediated increase of neutrophil numbers post viral infection (Fig. 4, H and I). Together, these results suggest that the microbiome-dependent immune cue(s) engage TLR2 to enhance neutrophil and type I IFN response in zebrafish post SVCV challenge, which culminated in restriction of viral infection.

### *C. somerae* mono-colonization recapitulates the antiviral effect of the intestinal microbiome

We next investigated whether specific bacterial taxa mediated the antiviral effect of the microbiome. We tested this idea by evaluating whether a single antibiotic was required or dispensable for the effect in increasing the susceptibility to viral infection. We feed adult zebrafish with antibiotics cocktail including neomycin (Neo), vancomycin (Vanco), ampicilin (Amp), and metronidazole (Metro) or with each antibiotic singly. Two additional single antibiotic treatments Clindamycin (CLI, an antibiotic especially efficient against anaerobic bacteria) and Nalidixic acid (NAL, an antibiotic predominantly targeting gram-negative bacteria) were also included. Zebrafish were fed for 2 weeks and challenged with SVCV through intraperitoneal injection. The mortality of zebrafish was consistently increased by antibiotics cocktail (Fig. 5, A and B). Interestingly, Metro and CLI treatment led to increased mortality similar to the cocktail group, suggesting that the two antibiotics depleted key bacterial taxa responsible for the antiviral function (Fig. 5, A and B). In contrast, Neo group showed similar mortality with control, suggesting that the bacterial taxa eliminated by Neo were dispensable (Fig. 5, A and B). Consistently, GF zebrafish colonized with microbiota from control and Neo group of fish showed lower viral infection compared with GF control, while colonization with microbiota from Metro and CLI group of fish had no effect (Fig. 5C). Therefore, we carried out 16S *r*RNA gene sequencing on the intestinal contents collected from control fish and those fed antibiotics cocktail, Metro, CLI, or Neo at 0 dpi. Amp, Vanco, and NAL treatment exhibited a partial suppressive effect in resistance against viral infection and thus were not included in the sequencing.

**Fig. 5.**
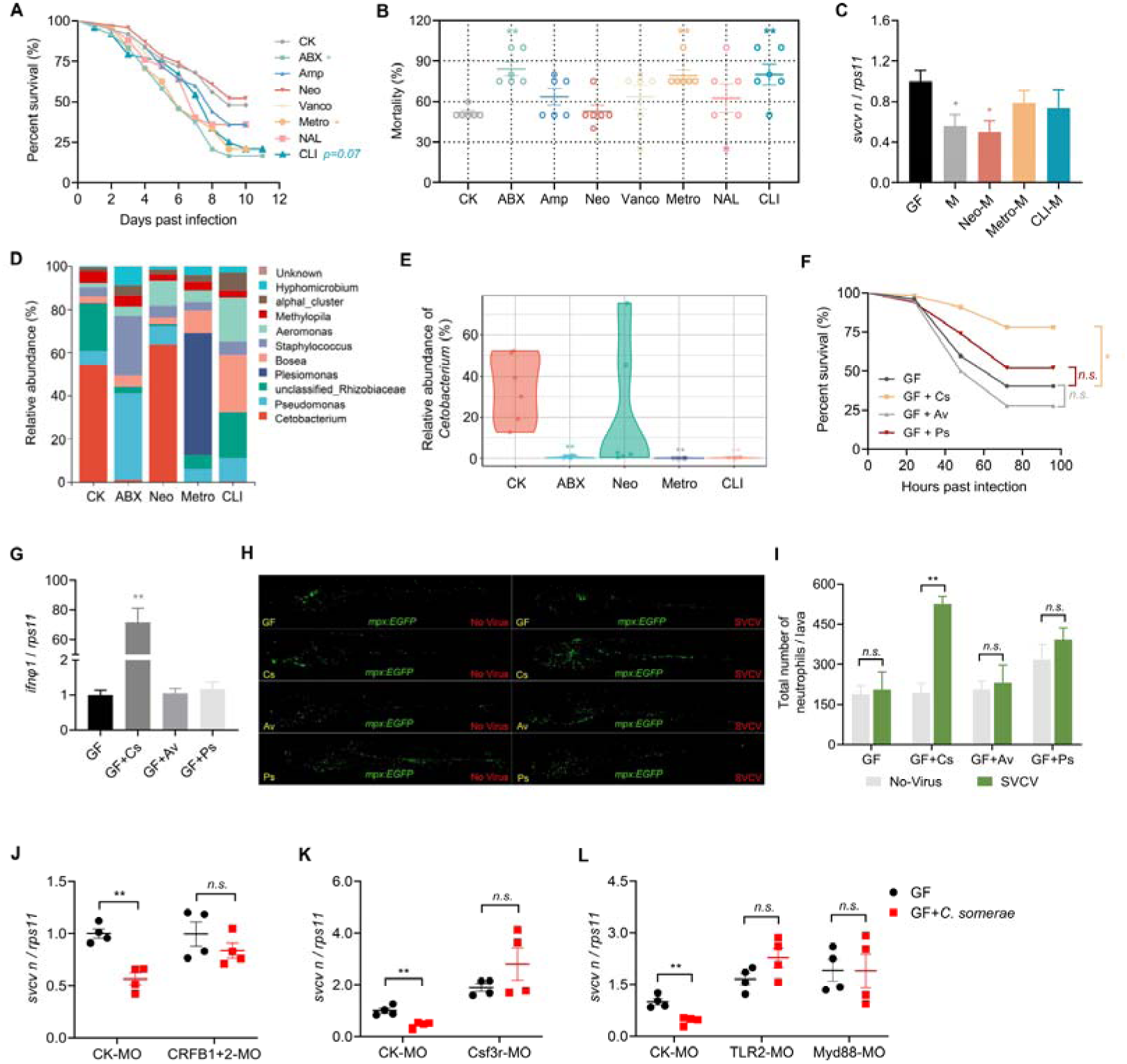
*C. somerae* recapitulates the antiviral effect of intestinal microbiome. (**A**-**B**) Effect of antibiotics cocktail or single antibiotic feeding on SVCV infection in adult zebrafish. (A) Survival curve (n=25), (B) Mortality (n=6). (**C**) Viral replication in GF zebrafish or GF zebrafish colonized with microbiota derived from adult zebrafish fed with control or antibiotic(s) diet. (n = 3, pool of 30 zebrafish larvae per sample). Mock, GF group; M, control microbiota; Neo-M, microbiota from zebrafish fed neomycin; Metro-M, microbiota from zebrafish fed metronidazole; CLI-M, microbiota from zebrafish fed clindamycin. (**D**) The composition of intestinal microbiota of adult zebrafish fed control or antibiotic(s) diet. (n =6, pool of 6 zebrafish per sample). (**E**) The relative abundance of *Cetobacterium* in intestinal microbiota of adult zebrafish fed control or antibiotic(s) diet. (n =6, pool of 6 zebrafish per sample). (**F**) Survival curve of GF zebrafish or GF zebrafish mono-colonized with *C. somerae* (GF+CS), *Aeromonas veronii* (GF+AV), or *Plesiomonas shigelloides*. (GF+PS) following SVCV infection (n = 60). (**G**) IFNΦ1 expression of GF zebrafish or GF zebrafish mono-colonized with indicated commensal bacterium. Expression was detected at 48 hpi. (n = 4, pool of 30 zebrafish larvae per sample). (**H**-**I**) Neutrophils response in mock or SVCV-infected GF *Tg* (*mpx:EGFP*) zebrafish or GF counterparts mono-colonized with indicated commensal bacterium. (H) Confocal imaging of transgenic zebrafish, (I) Neutrophil numbers (n=3). Samples were collected for imaging at 48 hpi. Scale bar, 500 μm. (**J**-**L**) Effect of type I IFN receptors knockdown (CRFB1+CRFB2 MO) (J), depletion of neutrophils (Csf3r MO) (K), and TLR2 and Myd88 knockdown (L) on SVCV infection in GF zebrafish or GF counterparts mono-colonized with *C. somerae* (n = 4, pool of 30 zebrafish larvae per sample). Viral replication was detected at 48 hpi. (A and F) log-rank test; (B, C, E, G) one-way ANOVA followed by Dunnett’s multiple comparisons test; (I-L) unpaired *t* test. \**p*< 0.05, \*\**p*< 0.01, *n.s.*, not significant.

The 16S *r*RNA gene sequencing result clearly showed that the abundance of *Cetobacterium* was correlated with the protective effect of the microbiota. Indeed, *Cetobacterium* was depleted in ABX, Metro and CLI groups with high mortality, while its abundance was high in the control and Neo groups (Fig. 5, D and E, fig. S7). We mono-colonized germ-free zebrafish with *C. somerae* isolated from the intestine of zebrafish and challenged the gnotobiotic fish with SVCV. Remarkably, colonization of *C. somerae* reduced SVCV infection compared with GF control (Fig. 5F). In comparison, mono-colonization of *Aeromonas veronii* and *Plesiomonas shigelloides*, two other commensal bacterial strains isolated from the intestine of zebrafish, did not inhibit viral infection (Fig. 5F), thus controlling for any non-specific effects due to bacterial colonization. Furthermore, we tested the effect of *C. somerae* in adult zebrafish. Results showed that *C. somerae* reduced the mortality of zebrafish after SVCV infection (fig. S8).

We hypothesized that *Cetobacterium* inhibited SVCV infection by stimulating neutrophil-dependent type I IFN response. Indeed, *C. somerae* colonization enhanced neutrophil response and IFNΦ1 expression versus GF control following SVCV infection, while *Aeromonas veronii* and *Plesiomonas shigelloides* did not have the effects (Fig. 5, G to I). Consistently, the antiviral function of *C. somerae* was abrogated by CRFB1+2 knockdown as well as Csf3r knockdown-mediated neutrophil depletion (Fig. 5, J and K). Moreover, we found that the protective effect of *C. somerae* required TLR2-Myd88 signaling, as the bacterium-mediated reduction of viral replication was blocked by TLR2 and Myd88 knockdown (Fig. 5L). Thus, *C. somerae* restricted SVCV infection by a mode that mirrored the microbiome, indicating a key role of this bacterial taxon in the microbiome-mediated function.

### The exopolysaccharides of *C. somerae* signals through TLR2 to mediate antiviral effect

To identify the bacterial components responsible for the antiviral effect, we treated GF zebrafish with cell free supernatant or cell lysate of *C. somerae* and then challenged them with SVCV (Fig. 6A). Both the cell free supernatant and cell lysate reduced viral infection (Fig. 6, B and C). Next, both the cell free supernatant and cell lysate were separated using a 3 kDa centrifugal filter. We tested the function of both the upper retentate (macromolecules) and the lower liquid filtrate (small-molecule compounds) in GF zebrafish. Intriguingly, treatment of GF zebrafish with the retentate rather than the filtrate reduced viral infection, indicating that the effector(s) are macromolecule(s) (Fig. 6, B and C). Furthermore, the antiviral function of both cell free supernatant and cell lysate was maintained following proteinase K treatment, suggesting that the effector(s) are not protein(s) and probably are polysaccharide(s) (Fig. 6, B and C).

**Fig. 6.**
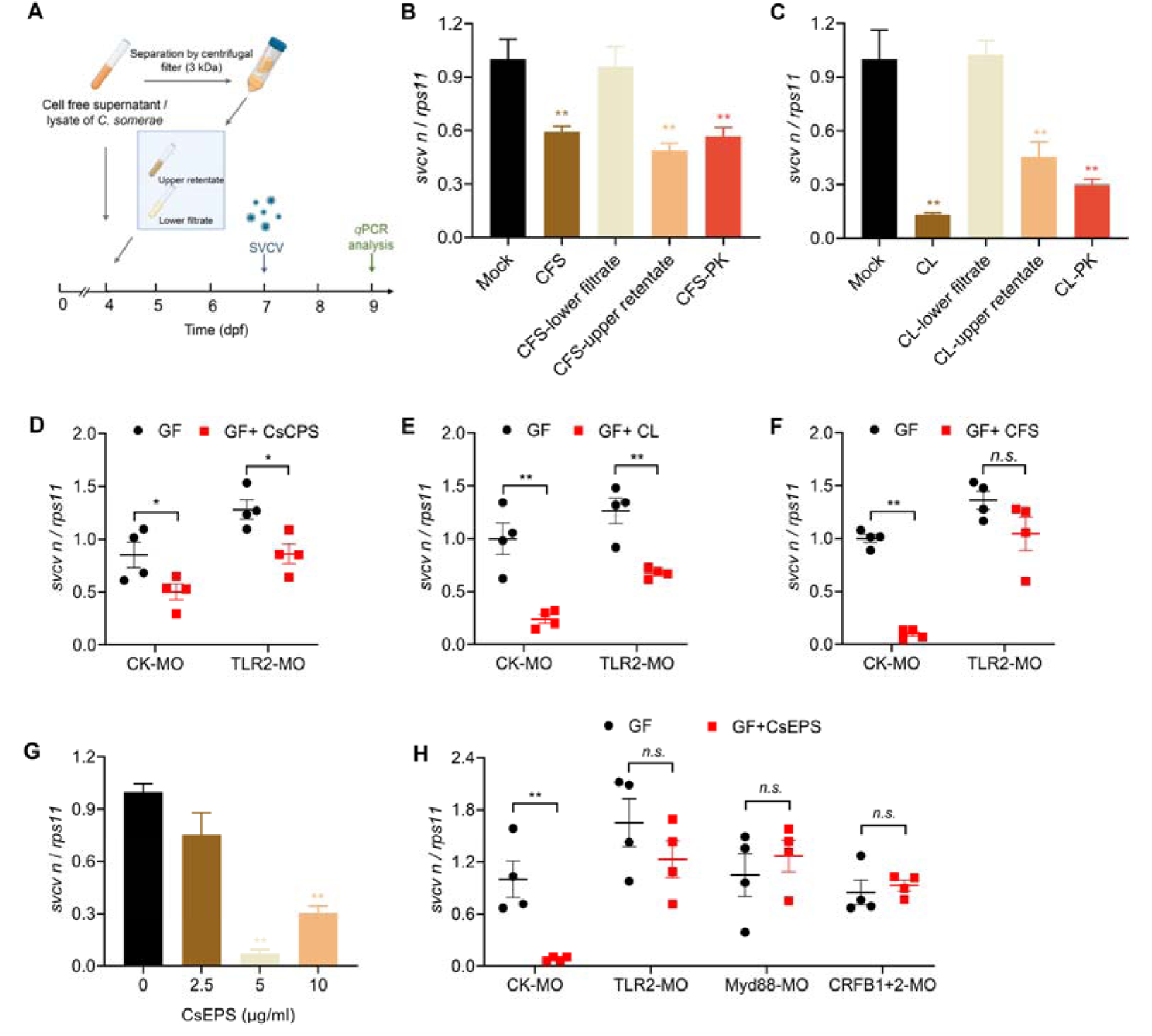
Exopolysaccharides of *C. somerae* signal through TLR2 to inhibit SVCV infection. (**A**) Schematic representation of the study design. (**B**-**C**) Nonprotein macromolecule(s) mediated the antiviral effect of *C. somerae*. The *C. somerae* culture suspension was separated into cell free supernatant (CFS) and bacterial cells by centrifugation. CFS and cell lysate (CL) were separated by a 3kDa filter, or treated with proteinase K. GF zebrafish were treated with different CFS or CL samples and subjected to SVCV infection. Viral replication was detected at 48 hpi. (n = 4, pool of 30 zebrafish larvae per sample). CFS, cell free supernatant; CL, cell lysate; “-lower filtrate”, 3kDa filtrate; “-upper retentate”, 3 kDa upper retentate; “-PK”, proteinase K treated CFS or CL samples. **(D)** Effect of morpholino-mediated TLR2 knockdown on the antiviral effect mediated by *C. somerae* CPS in GF zebrafish. Viral replication was detected at 48 hpi. (n = 4, pool of 30 zebrafish larvae per sample). (**E-F**) Effect of morpholino-mediated TLR2 knockdown on the antiviral effect mediated by *C. somerae* CL (E) or CFS (F) in GF zebrafish. Viral replication was detected at 48 hpi. (n = 4, pool of 30 zebrafish larvae per sample). (**G**) *C. somerae* exopolysaccharides (CsEPS) inhibited SVCV infection in GF zebrafish. GF zebrafish were treated with different doses of CsEPS and subjected to SVCV infection. Viral replication was detected at 48 hpi. (n = 4, pool of 30 zebrafish larvae per sample). (**H**) Effect of morpholino-mediated knockdown of TLR2, Myd88, and type I IFN receptors on the antiviral effect of CsEPS in GF zebrafish. GF zebrafish were treated with CsEPS at 5 μg/mL and subjected to SVCV infection. Viral replication was detected at 48 hpi. (n = 4, pool of 30 zebrafish larvae per sample). (B, C, G) One-way ANOVA followed by Dunnett’s multiple comparisons test; (D, E, F, H) unpaired *t* test. \**p*< 0.05, \*\**p*< 0.01, *n.s.*, not significant.

We observed that *C. somerae* contained a capsule structure by capsular staining (fig. S9A). Capsular polysaccharides (CPS) have been reported to mediate antiviral effect of commensal *Bacteroides* in mammals (*2*). We extracted the CPS from *C. somerae*, and found that *C. somerae* CPS (CsCPS) exhibited significant antiviral activity (fig. S9B). However, the antiviral effect of CPS was maintained in TLR2 morphant zebrafish, suggesting that CPS was not the key effector responsible for *C. somerae*-mediated function (Fig. 6D). We also excluded peptidoglycan as the effector by lysozyme treatment of cell lysate (fig. S10). Furthermore, we isolated LPS of *C. somerae* and found that the LPS exhibited no antiviral function (fig. S11). We then hypothesized that exopolysaccharides secreted in the supernatant were the main antiviral effector. Indeed, the antiviral effect of cell lysate was maintained in TLR2 morphant zebrafish, while the effect of cell free supernatant was abrogated by TLR2 knockdown (Fig. 6, E and F). We then extracted exopolysaccharides from cell free supernatant of *C. somerae* and found that *C. somerae* exopolysaccharides (CsEPS) had significant antiviral activity (Fig. 6G). Intriguingly, the antiviral effect of CsEPS was blocked by TRL2 and Myd88 knockdown (Fig. 6H), supporting that exopolysaccharides were the *C. somerae*-derived TLR2 ligand that played a key role in the *in vivo* antiviral function. The antiviral effect of CsEPS was abrogated in CRFB1+2 morphant zebrafish (Fig. 6H), which was also consistent with *C. somerae* mono-colonization result. Together, these results demonstrate that a single commensal bacterial molecule limits viral infection in a mode resembling the whole microbiota, suggesting a CsEPS-TLR2-type I IFN signaling axis in mediating the antiviral effect of microbiome in zebrafish.

## DISCUSSION

We have revealed a previously unknown role of the intestinal microbiome in promoting type I IFN antiviral response in zebrafish. Our results are consistent with previous reports of commensal regulation of type I IFN response in mammalian viral infection models (*8–11*), suggesting a conserved role of commensal bacteria in regulating type I IFNs and viral infection among vertebrates. Notably, type II and III IFNs were reported to be involved in the antiviral effect of microbiome in some studies (*7*, *13*). Type III IFNs have not been identified in teleost (*23*, *25*). Recently, type IV IFN has been identified in zebrafish and other vertebrates, which also possesses antiviral activity (*30*). However, we observed no difference in the expression of type II and IV IFNs between GF and conventionalized zebrafish after SVCV infection, excluding the involvement of other IFN types in the antiviral effect of microbiome. Previous studies demonstrated that the microbiome regulates type I IFN production under steady state conditions (*2*, *36*). However, in our analysis, differences in the expression of type I IFN genes were not observed between conventionalized zebrafish and the GF controls at steady state, probably due to that the *q*PCR method we used was not sensitive enough to detect the differences.

Our results showed that neutrophils mediated the antiviral effect of microbiome in zebrafish. Depletion of neutrophils also abrogated the microbiota-mediated stimulation of IFNΦ1 expression. Neutrophils are the main IFN-producing leukocyte in zebrafish upon Chikungunya virus (CHIKV) infection (*31*). In our study, we have no direct evidence to support that neutrophils were the IFN-producing cells after viral infection. It is also possible that neutrophils are not the main IFN producer, but stimulate the IFN production by other cells. We also observed that the expression of *mpx* (marker gene for neutrophils) was lower in the ABX group versus control at 1 and 2 days post SVCV infection or poly(I:C) treatment in adult zebrafish, while no difference was observed for the expression of *mpeg1* (marker gene for macrophages) (fig. S12). This suggested that the microbiome contributed to the activation of neutrophils, but not macrophages, in adult zebrafish post viral infection, which is consistent with the results in gnotobiotic zebrafish. Previous studies in mammalian model revealed that macrophages and pDCs were the effector immune cells that mediate the antiviral effect of microbiome (*13*, *37*). The discrepancy might be due to species-specific interaction of microbiome and immune cells that mediate the innate antiviral immunity.

Previous studies have suggested the contribution of microbe-associated molecular pattern (MAMPs) in priming of the antiviral immunity by intestinal microbiome. Ichinohe et al. found that rectal inoculation of agonists of TLR4, TLR3, TLR9, and to a lesser extent TLR2, rescued the antiviral immunity against influenza in ABX-treated mice (*38*). Zhang et al. reported that bacterial flagellin inhibited rotavirus infection in mice, which involved TLR5 and NOD-like receptor C4 (NLRC4) (*1*). However, these studies used canonical PRR agonists, and did not uncover the PRR signaling and identity of specific commensal microbes that mediate the priming of antiviral immunity by the microbiome. Diamond’s group demonstrated that restriction of CHIKV dissemination by the microbiome requires TLR7-Myd88 signaling in pDCs, but whether the identified bacterial species (*Clostridium scindens*) and its derived metabolite (the secondary bile acid deoxycholic acid) engage TLR7 to mediate the antiviral effect was not determined (*12*). In another study, Stefan et al. found that *Bacteroides fragilis* and its OM-associated polysaccharide A induce IFN-β via TLR4-TRIF signaling, which at least in part mediates the regulatory effect of microbiome on natural resistance to viral infection (*2*). The results in our study provided novel insights into the identity of specific commensal bacteria that primes the antiviral immunity as well as the underlying signaling pathway. We found that the microbiota-mediated antiviral effect relied on TLR2-Myd88 signaling, and further revealed that the commensal bacterium *C. somerae* played a major role in the TLR2-Myd88 dependent priming of the innate antiviral immunity via its exopolysaccharides. It is intriguing that TLR2 activation by a gram negative bacterium primed antiviral immunity, as TLR2 is a canonical PRR for peptidoglycan. The sequencing result showed considerable abundance of *Staphylococcus* in the microbiota of zebrafish, but its relative abundance was not correlated with the antiviral effect of the microbiome. It might be that the *Cetobacterium* exopolysaccharides is a more potent agonist of TLR2 in zebrafish compared with peptidoglycan, or that *Cetobacterium* produces relatively higher amount of exopolysaccharides, which resulted in its dictating effect in priming TLR2-mediated antiviral immunity. Consistent with our results, previous study in gnotobiotic zebrafish demonstrated that TLR2 was essential for the regulation of microbiota on *myd88* transcription and intestinal immunity (*17*), which highlighted the key role of TLR2 in mediating the immunomodulatory effect of microbiome in zebrafish.

Co-evolution brings about a triangular relationship of host, microbiome, and viruses (*3*, *4*). Differential mechanisms have been reported to mediate the antiviral effect of microbiome, including both immune-dependent or –independent actions (39). In particular, priming of type I IFN response was implicated as the underlying mechanism in a number of studies (*8–11*). However, few studies have revealed immune cells and signaling pathways that link the microbiome to type I IFN response. Our data reveal that the microbiome primes type I IFN response through a pathway that requires neutrophils and TLR2-Myd88 signaling. Although the mechanisms involve different immune cells and TLRs compared with those in mammalian model (*2*, *13*), the underlying consistency suggests an evolutionally conserved function of the microbiome in regulation of type I IFN response and thus natural resistance against viral infection in lower vertebrates. Considering the advantages of zebrafish as an animal model, the conserved function suggests that gnotobiotic zebrafish can be used as a model to study the molecular foundations of the triangular interactions of host, microbiome, and virus, or as a screening platform for potential microbiome-derived antiviral molecules.

## MATERIALS AND METHODS

### Zebrafish husbandry

This study was approved by the Feed Research Institute of the Chinese Academy of Agricultural Sciences Animal Care Committee under the auspices of the China Council for Animal Care (Assurance No. 2015-AF-FRI-CAAS-003). Zebrafish were raised in a circulating water system under a 14 h/10 h light/dark cycle at 28°C and fed two times per day as previously described (*40*). Experiments were performed using the wild-type AB zebrafish strain. When necessary, transgenic line *Tg* (*mpx*:*EGFP*) and *Tg* (*mpeg1*:*EGFP*) were used for immune cell visualization.

### Antibiotics feeding

For antibiotics feeding experiment, adult AB zebrafish (two months old) were fed with antibiotic cocktail-supplemented feed for 14 days. The antibiotic cocktail contains ampicillin (Amp, 0.5 mg/kg), metronidazole (Metro, 0.5 mg/kg), neomycin (Neo, 0.5 mg/kg), and vancomycin (Van, 0.25 mg/kg). For single antibiotic feeding, zebrafish were fed with a single antibiotic of the cocktail or clindamycin (CLI, 0.11 mg/kg), nalidixicacid (NAL, 0.33 mg/kg). Antibiotics were purchased from Sigma-Aldric. Ingredients and proximate composition of diets for zebrafish are shown in table S1.

### Germ-free (GF) zebrafish and treatment

Germ-free zebrafish were prepared following established protocols as described previously (*41*). Zebrafish larvae were hatched from their chorions at 3 dpf with 30 fish per bottle. The transfer of gut microbiota from adult zebrafish to germ-free zebrafish was performed as previously describe (*42*). For mono-association, *C. somerae* XMX-1was incubated anaerobically in Gifu Anaerobic Medium (GAM) broth at 28°C for 12 h, while *Aeromonas veronii* XMX-5 and *Plesiomonas shigelloides* were cultured in Luria-Bertani (LB) broth at 37°C for 18 h. Then, the bacterial cells were harvested by centrifugation (7000 g for 10 min at 4°C). The collected bacterial cells were washed three times with distilled PBS. GF zebrafish were colonized at 4 dpf with a single bacterium at a final concentration of 10^5^ CFU/mL.

### Viral infection

Epithelioma papulosumcyprini (EPC) cells and SVCV (ATCC: VR-1390) were presented by professor Jun-Fa Yuan (Huazhong agricultural university, Wuhan, Hubei, China). SVCV was propagated in EPC cells as previously described (*43*). TCID_50_/mL was calculated according to the Reed and Muench method (*44*).

Adult zebrafish (two months old) was acclimatized to 22°C and were i.p. injected with SVCV (2.5 × 10^5^ 50% tissue culture-infective dose [TCID_50_]/ml) at 10 μL/individual. The liver, spleen, and kidney of zebrafish were collected 24 h and 48 h after injection. Mortality was recorded for 10 days. Germ-free, conventional, conventionalized, or mono-associated zebrafish were infected with SVCV at 7 dpf by bath immersion at a concentration of 10^6^ TCID_50_/mL. The infection was conducted at 25°C. Larval zebrafish were harvested at 9 dpf for *q*PCR or fluorescence analysis. Mortality was recorded for 96 hours after infection when necessary.

### Confocal microscopy

Images were captured by using a confocal ZEISS LSM 980 with the Airyscan2 super-resolution mode (Tsinghua University, China). For live-imaging, zebrafish larvae were anesthetized with 0.16 mg/mL of tricaine in embryo medium and mounted in 1.2% low-melting agarose on a cover slip with extra embryo medium sealed inside vacuum grease to prevent evaporation. Imaging was performed on a single z plane at 0.5 s intervals for 20-30 min. The images of *Tg* (*mpx*:*EGFP*) zebrafish were processed and reconstructed by Imaris 9.0.1 64-bit version (Bitplane, Switzerland), and neutrophils were enumerated by the same software. For images of *Tg* (*mpeg1*:*EGFP*) zebrafish, fluorescence intensity was calculated through Image J.Average flourescence intensity = The total fluorescence intensity in this area /Regional area.

### Gut microbiota analysis

At the end of the 2□week feeding trial, the gut contents of adult zebrafish were collected 4 h post the last feeding. The gut contents were collected under aseptic conditions. Each gut content sample was pooled from 6 fish. Bacteria DNA was extracted using the Fast DNA SPIN Kit for Soil (MP Biomedicals). The 16S V3−V4 region was amplified by using the primers as follows, 338F: 5′□ACTCCTACGGGAGGCAGCA□3′ and 806R: 5′□GGACTACHVGGGTWTCTAAT□3′. 16S *r*RNA gene sequencing was conducted at Biomarker Technologies using the illumina novaseq. 6000 platform (Illumina). Data analysis was performed using BMKCloud (www.biocloud.net). Microbiota sequencing data in this study are available from the National Center for Biotechnology Information (NCBI) under accession number PRJNA1115092.

### Morpholino knockdown

The MOs used in this study are all vivo-morpholino. Vivo-morpholino oligonucleotides (MO) were designed and synthesized by Gene-Tools (Philomath, OR). The sequences of MO used in this study are listed in table S2. MO was added to GZM at 4 dpf at 50-100 nM, except for Spi1b MOs, Csf3r MO, and Irf8 MO, which was added at 1 dpf. For Spi1b, two targeting MOs were simultaneously added to GZM at 1 dpf at concentrations indicated in table S2. For all the MOs treatment, zebrafish larvae were treated with MO throughout the following experimental period.

### Preparation and fractionation of cell free supernatant and cell lysate of *C. somerae*

*C. somerae* XMX-1 was grown in GAM broth at 28°C for 12 h. Cell free supernatant (CFS) was obtained by centrifugation (7000 g, 10 min) and filtration through a 0.22 µm filter (Millipore, Darmstadt, Germany). The bacterial cell pellets were washed three times with phosphate buffer (50 mM, pH 7.0) and resuspended in same volume of phosphate buffer. The filtered samples were incubated for 10 min on ice and were then subjected to sonication (300 W, with a pulse on/off ratio of 1 second on and 1 second off, for a total duration of 10 minutes). Sonicated bacterial sample was centrifuged at 7000 g for 5 min, and the supernatant was filtered aseptically through a 0.22 µm filter to obtain cell lysate of *C. somerae*. The cell free supernatant or cell lysate was separated by using a 3 kDa cutoff filter. Both the upper retentate and the lower filtrate were collected for bioassay. For experiments using proteinase K, CFS or cell lysate was incubated with protease K at 65°C for 2 h before inactivating the enzyme at 99°C for 30 min. All relevant bacterial samples were added to GF zebrafish at 4 dpf by immersion.

### Purification of polysaccharides from *Cetobacterium somerae*

Capsular polysaccharides were extracted from bacterial cell pellets of *C. somerae* by using a commercial CPS extraction kit (Genmed Scientifics Inc., USA). Bacterial exopolysaccharides purification was carried out according to the instruction manual of the commercial kit (Genmed Scientifics Inc., USA). Briefly, bacterial cells were swabbed from GAM agar plates, resuspended in salt-based media, and incubated for four hours at 25°C. Bacterial culture were centrifuged at 10000 g for 20 minutes. The supernatant was recovered and was then filtered using a 0.22 µm filter and concentrated through an Amicon Ultra centrifuge filter with a 100 kDa molecular weight cutoff. The filtered sample was treated with DNase, RNase and proteinase K, respectively. Samples were extracted with Tris-saturated phenol-chloroform and precipitated by cold-ethanol.

### Western blotting

Larval zebrafish were homogenized in ice-cold HBSS buffer mixed with 1 mM PMSF and phosphatase inhibitors. Equivalent amounts of total protein were loaded into a 12% SDS-PAGE for electrophoresis and then transferred into a PVDF membrane (Millipore, USA). After blocking nonspecific binding with 5% skimmed milk in TBST, the PVDF membrane was incubated with primary antibodies, i.e., antibodies against anti-GAPDH (CST, 2118, 1:2000), anti-TLR2 (CST, 12276, 1:1000). Blots were imaged on a ChemiDoc (SynGene) using chemiluminescence detection with ECL western blotting substrate (Thermo Scientific, 34095).

### Quantitative PCR analysis

Total RNA was isolated from larval zebrafish or liver /spleen /kidney tissues of adult zebrafish with Trizol reagent (TaKaRa, Tokyo, Japan) following the manufacturer’s protocol. The extracted RNA was re-suspended in 30 μl RNase-free water then quantified with a BioTek Synergy™2 Multi-detection Microplate Reader (BioTek Instruments, Winooski, VT) and agarose gel electrophoresis. One microgram of total RNA was used for reverse transcription with Revert Aid™ Reverse Transcriptase (TaKaRa, Tokyo, Japan) according to the manufacturer’s instructions. The synthesized cDNA was stored at –20°C. Experimental methods about *q*PCR reaction were conducted as previously described (45). The primers used in the experiment were listed in table S3. *Ribosomal protein s11* gene (*rps11*) was used as the internal reference gene, and the data were statistically analyzed by 2^-ΔΔCT^ method.

### Statistical analysis

All data were performed using GraphPad Prism 8 software (GraphPad Software Inc. CA, USA). All data were expressed as mean ± SEM. Data involving more than two groups were analyzed using one-way ANOVA followed by Dunnett’s multiple comparison test. Comparisons between the two groups were analyzed using the unpaired Student’s *t* test. For the survival experiments, Kaplan–Meier survival curves were constructed and analyzed with the log-rank (Mantel–Cox) test. Statistical significance was denoted in figures as \**p*□≤□0.05, \*\**p*□≤□0.01, *n.s.*, not significant.

## ACKNOWLEDGMENTS

We are grateful to Dr. Jun-Fa Yuan for sharing the virus, and Bing-Yu Liu for helping with fluorescence microscopy.

## AUTHOR CONTRIBUTIONS

C. R. and Z. Z. designed the experiments; H. L. conducted the experiment and analyzed the data; M. L., J. C., W. Z., D. X., Q. D., Y. Y. and Z. Z. helped with the experiments or provided the reagents; C. R. and H. L. prepared the manuscript.

## Supplementary Materials

**PDF file includes:**

Tables S1, S2 to S3

Figs. S1 to S12

