## Supplemental Tables and Figures for "The intestinal microbiome and *Cetobacterium somerae* inhibit viral infection through TLR2-type I IFN signaling axis in zebrafish"

**SUPPLEMENTARY TABLES**

**Table S1. Ingredients and proximate composition of diets for zebrafish (dry matter, g/kg diet).**

| **Ingredient**  **(g / 100g diet)** | **Control** | **ABX** | **Ampicillin** | **Neomycin** | **Vancomycin** | **Metronidazole** | **nalidixic** | **clindamycin** |
| --- | --- | --- | --- | --- | --- | --- | --- | --- |
| Casein | 40 | 40 | 40 | 40 | 40 | 40 | 40 | 40 |
| Gelatin | 10 | 10 | 10 | 10 | 10 | 10 | 10 | 10 |
| Dextrin | 28 | 28 | 28 | 28 | 28 | 28 | 28 | 28 |
| **Ampicillin** | **0** | **0.5** | **0.5** | **0** | **0** | **0** | **0** | **0** |
| **Neomycin** | **0** | **0.5** | **0** | **0.5** | **0** | **0** | **0** | **0** |
| **Vancomycin** | **0** | **0.25** | **0** | **0** | **0.25** | **0** | **0** | **0** |
| **Metronidazole** | **0** | **0.5** | **0** | **0** | **0** | **0.5** | **0** | **0** |
| **nalidixic** | **0** | **0** | **0** | **0** | **0** | **0** | **0.332** | **0** |
| **clindamycin** | **0** | **0** | **0** | **0** | **0** | **0** | **0** | **0.1116** |
| Soybean oil | 6 | 6 | 6 | 6 | 6 | 6 | 6 | 6 |
| Lysine | 0.33 | 0.33 | 0.33 | 0.33 | 0.33 | 0.33 | 0.33 | 0.33 |
| VC phosphate ester | 0.1 | 0.1 | 0.1 | 0.1 | 0.1 | 0.1 | 0.1 | 0.1 |
| Vitamin mix^1^ | 0.4 | 0.4 | 0.4 | 0.4 | 0.4 | 0.4 | 0.4 | 0.4 |
| Mineralmix^2^ | 0.4 | 0.4 | 0.4 | 0.4 | 0.4 | 0.4 | 0.4 | 0.4 |
| CaH_2_PO_4_ | 2 | 2 | 2 | 2 | 2 | 2 | 2 | 2 |
| Choline chloride | 0.2 | 0.2 | 0.2 | 0.2 | 0.2 | 0.2 | 0.2 | 0.2 |
| Sodium alginate | 2 | 2 | 2 | 2 | 2 | 2 | 2 | 2 |
| Microcrystalline cellulose | 4 | 4 | 4 | 4 | 4 | 4 | 4 | 4 |
| **Zeolite** | **6.57** | **5.99** | **6.07** | **6.07** | **6.32** | **6.07** | **6.24** | **6.46** |
| Total | 100 | 100 | 100 | 100 | 100 | 100 | 100 | 100 |

^1^Containing the following (g/kg vitamin premix): thiamine, 0.438; riboflavin, 0.632; pyridoxine⋅HCl, 0.908; *d*-pantothenic acid, 1.724; nicotinic acid, 4.583; biotin, 0.211; folic acid, 0.549; vitamin B-12, 0.001; inositol, 21.053; menadione sodium bisulfite, 0.889; retinyl acetate, 0.677; cholecalciferol, 0.116; *dl*-α-tocopherol-acetate, 12.632.

^2^Containing the following (g/kg mineral premix): CoCl_2_⋅6H_2_O, 0.074; CuSO_4_⋅5H_2_O, 2.5; FeSO_4_⋅7H_2_O, 73.2; NaCl, 40.0; MgSO_4_⋅7H_2_O, 284.0; MnSO_4_⋅H_2_O, 6.50; KI, 0.68; Na_2_SeO_3_, 0.10; ZnSO_4_⋅7H_2_O, 131.93; Cellulose, 501.09.

**Table S2. Morpholino oligonucleotides.**

| **Target** | **Blocking type** | **Dose (nM)** | **Sequence of Morpholinos (5’-3’)** | **Reference** |
| --- | --- | --- | --- | --- |
| CK | — | 50/100 | CCTCTTACCTCAGTTACAATTTATA | — |
| TLR2 | translation | 75 | AGTCATTGTTCCTACGAGTCTCATC | (46) |
| TLR3 | splice | 75 | TCATTAGATCCATATTTTCCTTCCT | (47) |
| TLR4ba | translation | 75 | GATGCTGCTGAGGTTTCTTCCCATG | (45) |
| TLR7 | translation | 75 | TCATGGTCTTCTCAGTCATCTGAAA | (47) |
| TLR9 | translation | 75 | TCAAGGACACCATTGGTCCAAACAT | (47) |
| Spi1b | translation | 75 | CCTCCATTCTGTACGGATGCAGCAT | (48) |
| Spi1b | splice | 7 | GGTCTTTCTCCTTACCATGCTCTCC | (48) |
| Csf3r | translation | 50 | AAGCACAAGCGAGACGGATGCCAT | This study |
| Irf8 | translation | 50 | GCCCGAGTTCATCTTGTAGACCTTT | This study |
| Myd88 | splice | 50 | ATATCCACAAAGCAACATGCCTTTT | This study |
| CRFB1 | splice | 50 | CGCCAAGATCATACCTGTAAAGTAA | (49) |
| CRFB2 | splice | 50 | AGTTTGTTTTCTCACCTCTGTTCCA | This study |

**Table S3. Primer sequences for *q*PCR analysis.**

| **Gene** | **Nucleotide sequence of primers (5’-3’)** |
| --- | --- |
| *rps11* | F: ACAGAAATGCCCCTTCACTG |
|  | R: GCCTCTTCTCAAAACGGTTG |
| *svcvn* | F: TGAGGTGAGTGCTGAGGATG |
|  | R: CCATCAGCAAAGTCCGGTAT |
| *ifnφ1* | F: GAGCACATGAACTCGGTGAA |
|  | R: TGCGTATCTTGCCACACATT |
| *ifnφ2* | F: CCTCTTTGCCAACGACAGTT |
|  | R: CGGTTCCTTGAGCTCTCATC |
| *ifnφ3* | F: TTCTGCTTTGTGCAGGTTTG |
|  | R: GGTATAGAAACGCGGTCGTC |
| *ifnγ* | F: TGCACACCCCATCTTCCTGCGAA |
|  | R: GTGTTGCTTCTCTATAGACACGCTT |
| *ifnγrel* | F: TCAGACAACCAGCGCATACAGAT |
|  | R: TCATAGATGCTCAGCATTCCTTCA |
| *ifnυ* | F: CAACTGTGCGTTTGACTATGAT |
|  | R: CTCATTTTCAACACCGACCGAG |
| *crhb1* | F: CATCATACACTGGCCGTCTT |
|  | R: TGTCTGAAGTTGTGAGACCATATT |
| *crhb2* | F: TGGAACAGAGTTTACTGTGGATAAG |
|  | R: GTGGGCTTCGATCAATGATACT |
| *mpx* | F: GGGGCAGAAGAAGAAAGTCC |
|  | R: CCTTGCTAAACTCTCATCTC |
| *mpeg1* | F: CGATTGTGCCGCAAAAACT |
|  | R: GATTCATCACTCTGCCCATGTC |
| *myd88* | F: CCTGAAGCTTTGTGTGTTTGAC |
|  | R: TGACCACCACCATCCTCTTA |

**SUPPLEMENTARY FIGURES**

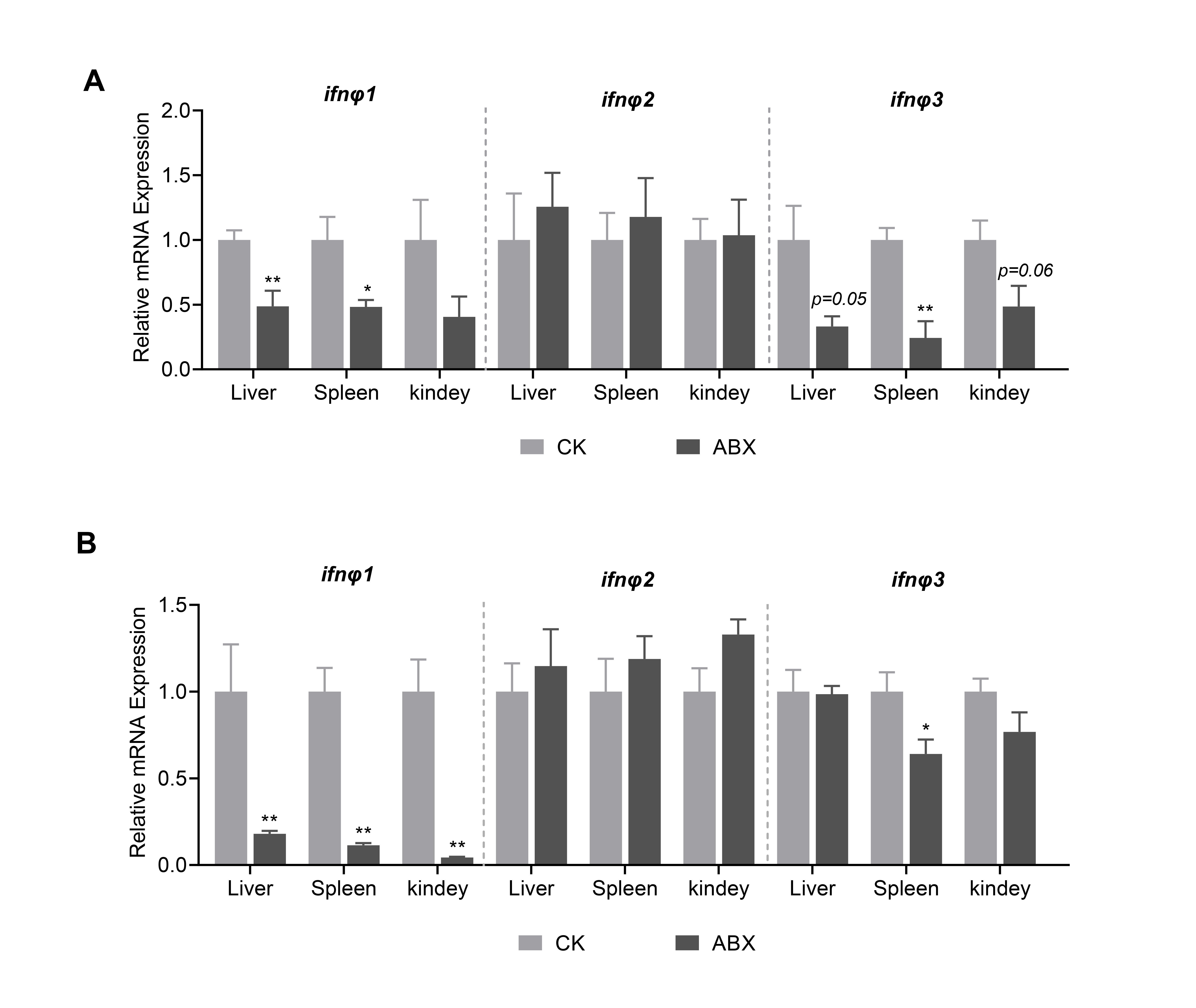

**Fig. S1. Intestinal microbiome depletion impairs type I IFN response to poly(I:C) treatment in adult zebrafish.** Expression of IFNΦ1, IFNΦ2, and IFNΦ3 in the liver, spleen, and kidney of adult zebrafish 1 (**A**) and 2 (**B**) days post poly(I:C) treatment (n = 4, pool of 6 fish per sample).

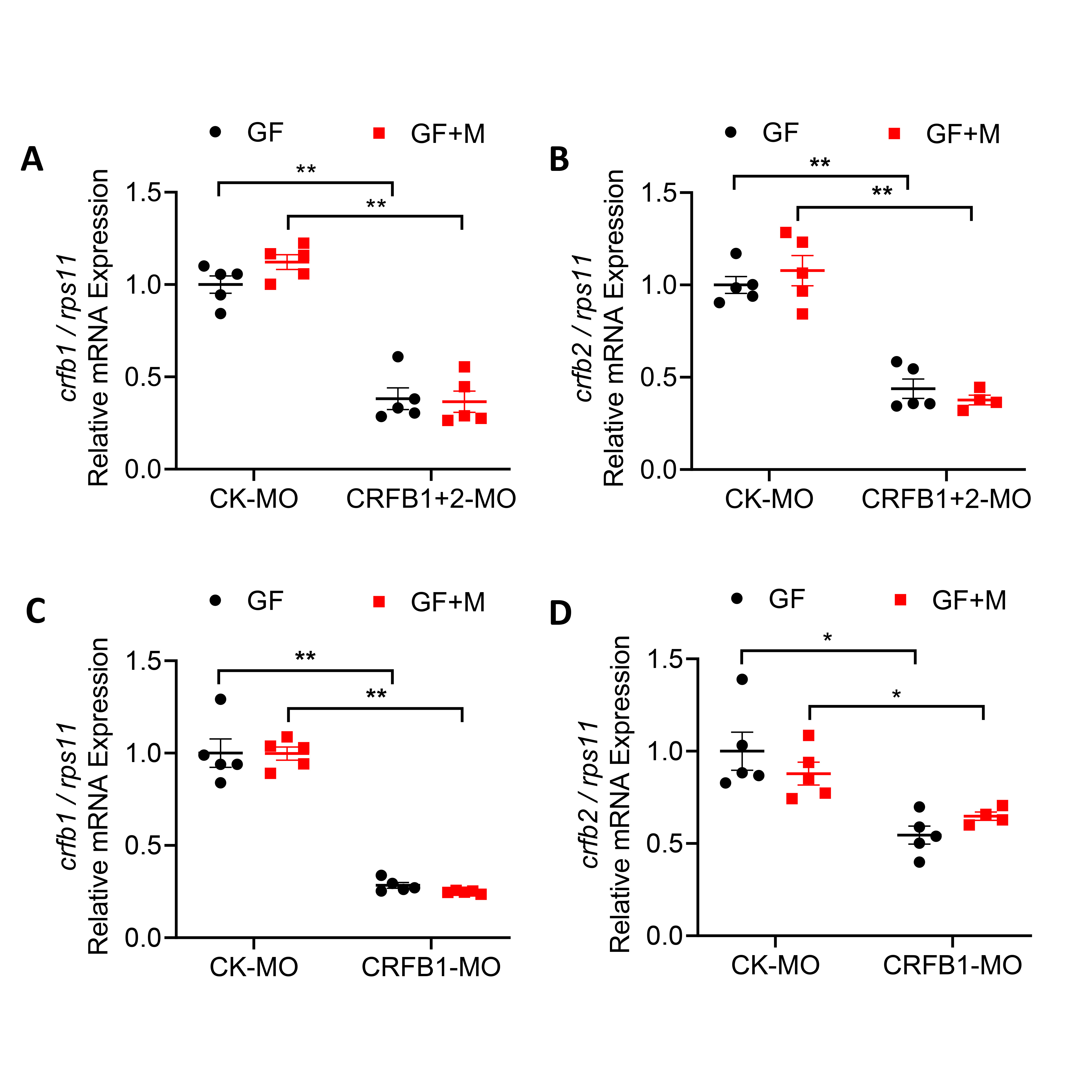

**Fig. S2. Knockdown efficiency of CRFB1+2 morpholino.** (**A**-**B**) GF and conventionalized zebrafish were treated with control morpholino (CK-MO) or a mixture of CRFB1 and CRFB2 morpholino (CRFB1+2 MO). Knockdown efficiency of CRFB1 (A) and CRFB2 (B) was detected by *q*PCR analysis of properly spliced transcripts (n = 5, pool of 30 zebrafish larvae per sample). (**C**-**D**) GF and conventionalized zebrafish were treated with control morpholino (CK-MO), CRFB1 morpholino (CRFB1-MO), or CRFB2 morpholino (CRFB2-MO). Knockdown efficiency of CRFB1 (C) and CRFB2 (D) was detected by *q*PCR analysis of properly spliced transcripts (n = 5, pool of 30 zebrafish larvae per sample).

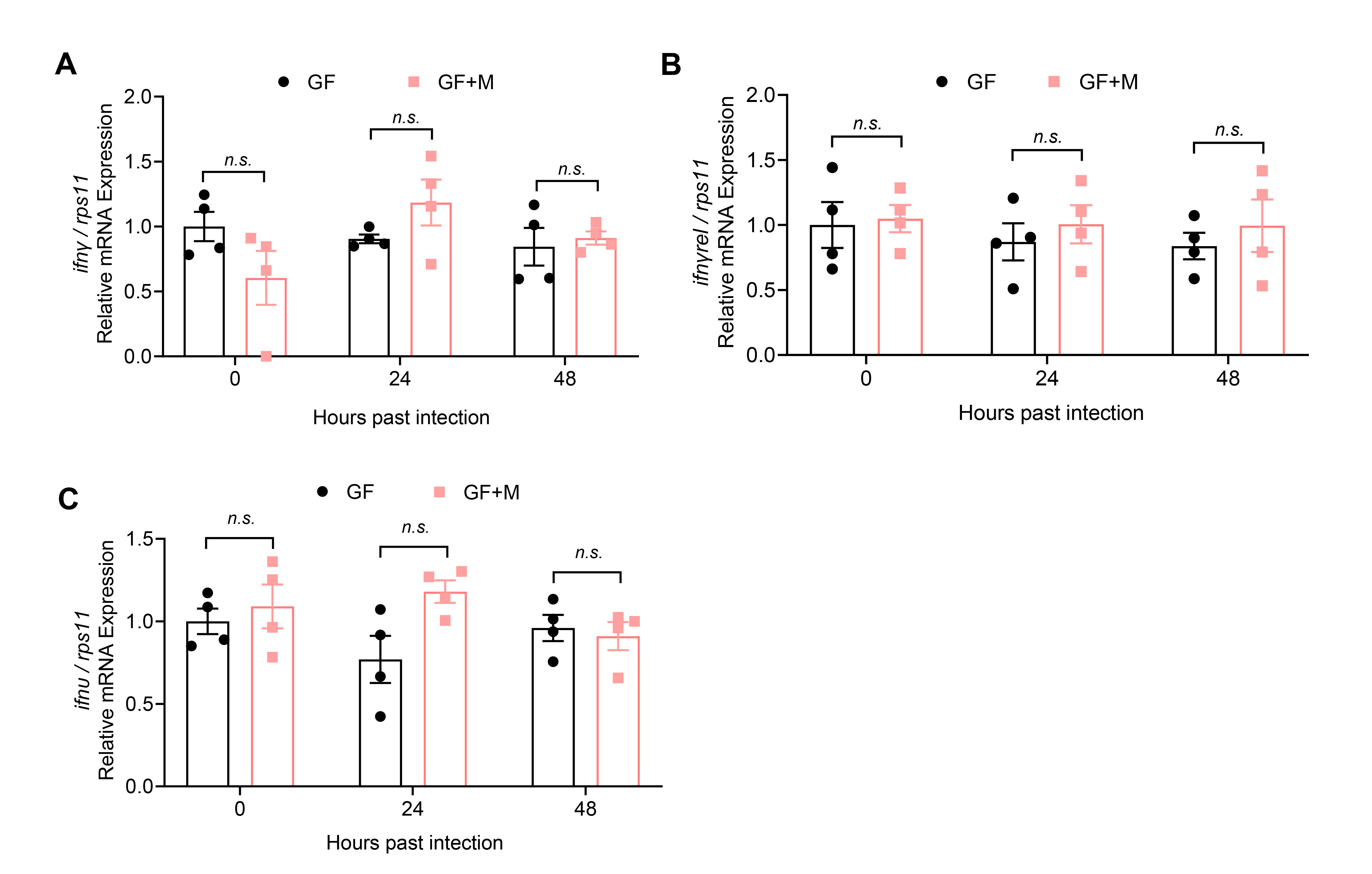

**Fig. S3. Expression of type II and type IV interferons after SVCV infection.** The expression of *ifnγ* (**A**), *ifnγrel* (**B**), and *ifnυ* (**C**) in GF or conventionalized zebrafish was detected at 0, 24, 48 hpi (n = 4, pool of 30 zebrafish larvae per sample).

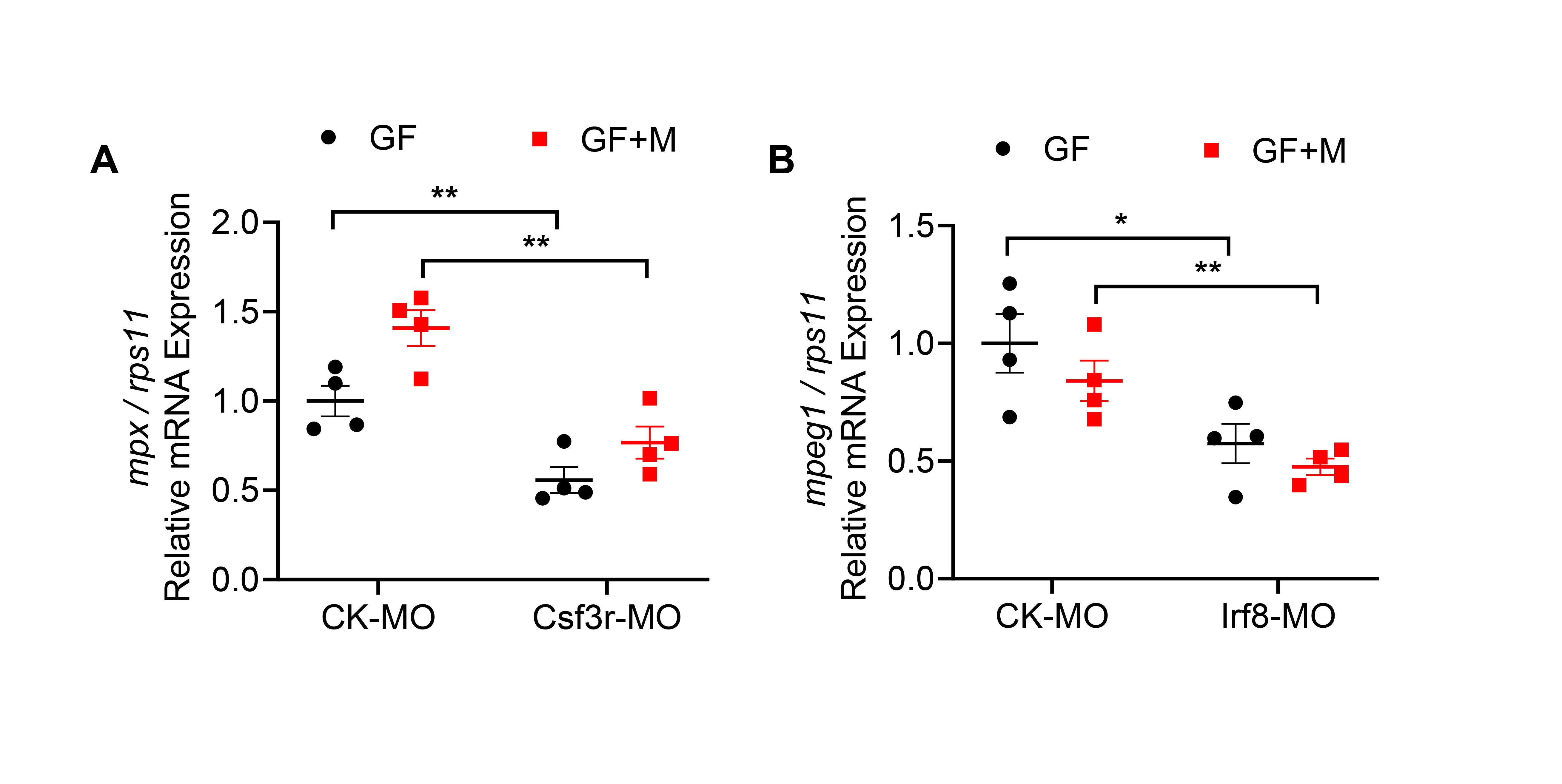

**Fig. S4. Depletion of neutrophils and macrophages by gene knockdown.** Effect of Csf3r MO (**A**) and Irf8 MO (**B**) on the expression of *mpx* (marker gene for neutrophil) (A) and *mpeg1* (marker gene for macrophage) (B) in GF or conventionalized zebrafish. (n = 4, pool of 30 zebrafish larvae per sample).

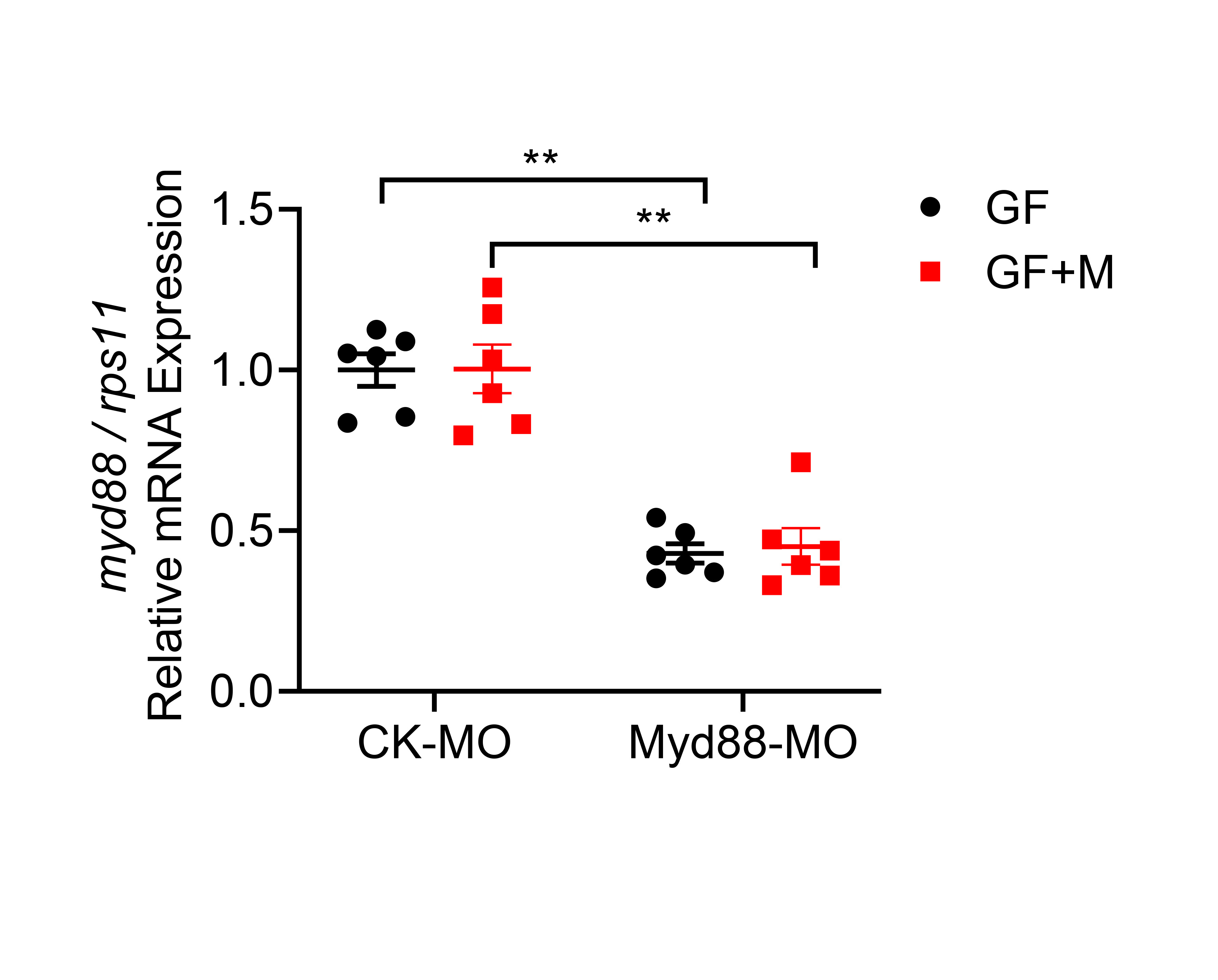

**Fig. S5. Knockdown efficiency of Myd88 morpholino.** Knockdown efficiency of Myd88 was detected by *q*PCR analysis of properly spliced transcripts (n = 6, pool of 30 zebrafish larvae per sample).

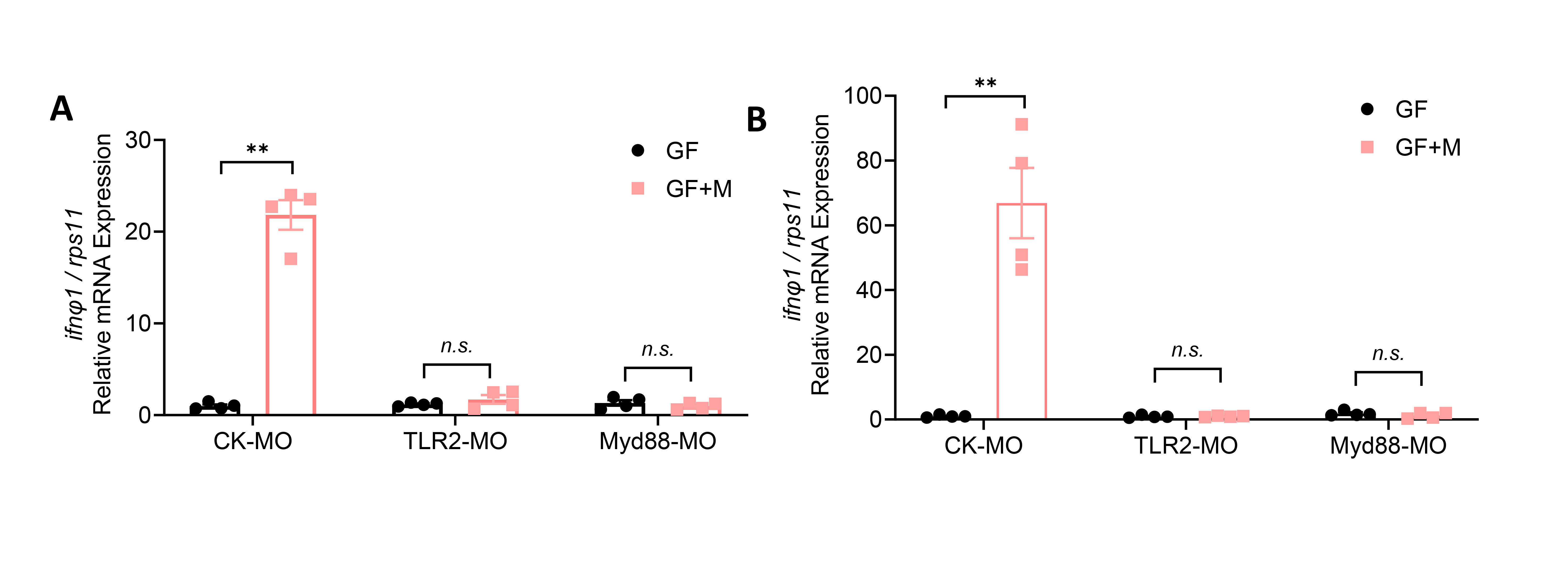

**Fig. S6. Effect of TLR2 and Myd88 knockdown on IFNΦ1 expression in GF or conventionalized zebrafish.** At 24 h (**A**) and 48 h (**B**) after poly(I:C) treatment (n=6, pool of 30 zebrafish larvae per sample).

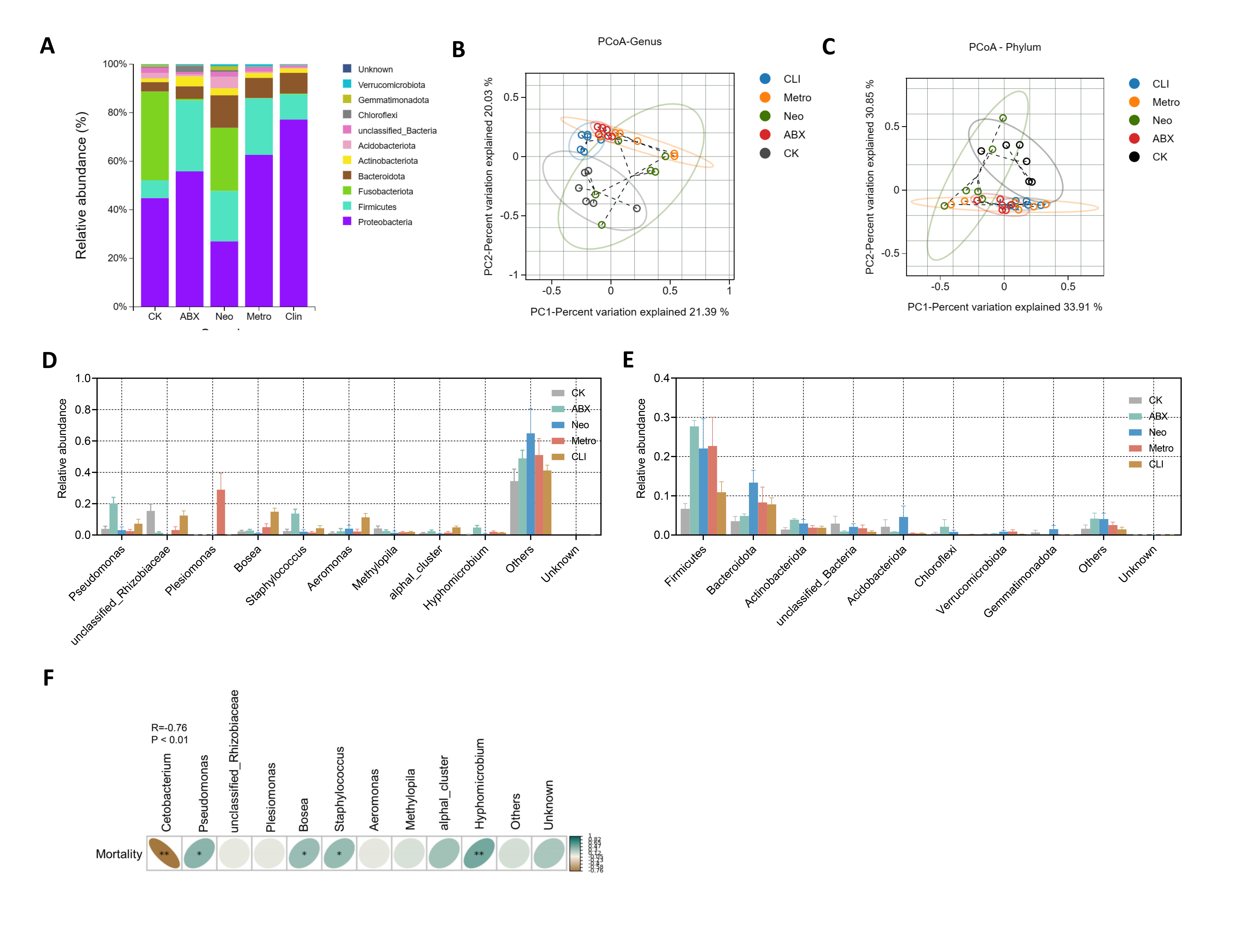

**Fig. S7. The composition of intestinal microbiome of adult zebrafish fed control or antibiotic(s) diet.** (**A**) Relative abundance of bacterial phyla. (**B**-**C**) Principal coordinate analysis (PCoA) of all samples by weighted UniFrac distance at the genus (C) and phylum (D) level (n =6, pool of 6 fish per sample)*.* (**D**-**E**) The relative abundance of specific taxa at genus (D) and phylum (E) level among groups. (**F**) Correlation analysis between the mortality of zebrafish and the relative abundance of genuses. (n =6, pool of 6 fish per sample).

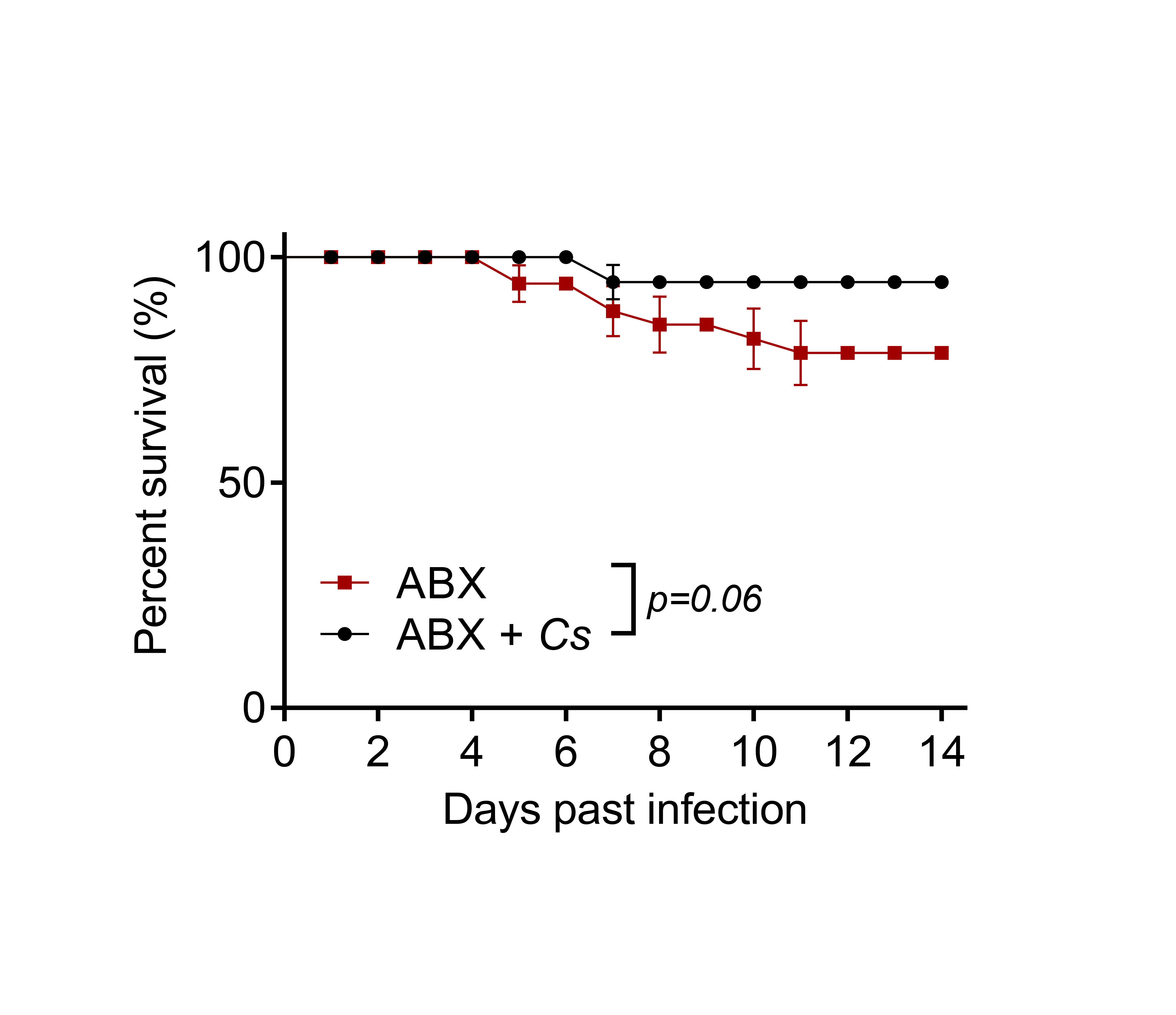

**Fig. S8. *C. somerae* inhibits SVCV infection in adult zebrafish.** Adult zebrafish were fed with antibiotics cocktail (ABX) for two weeks and were treated with PBS (mock) or *C. somerae* by immersion (ABX+Cs). Survival curve of zebrafish was monitored after SVCV infection. Survival was analysis by log-rank test (n = 41).

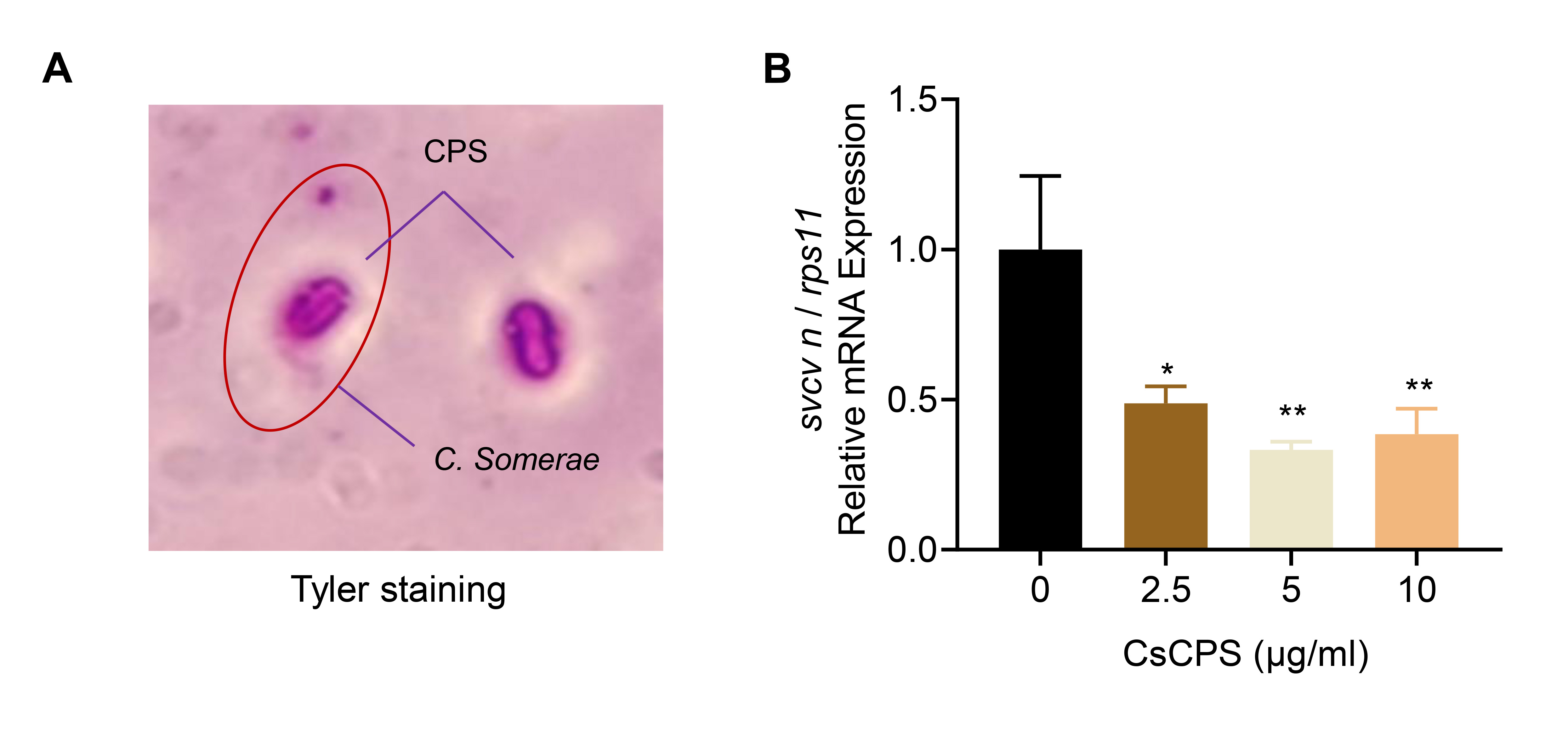

**Fig. S9. Antiviral activity of capsule polysaccharides (CPS) from *C. somerae*.** (**A**) Tyler staining forcapsule of *C. somerae*. (**B**) *C. somerae* CPS inhibited SVCV infection in GF zebrafish. GF zebrafish were treated with different doses of CPS and subjected to SVCV infection. Viral replication was detected at 48 hpi. (n = 4, pool of 30 zebrafish larvae per sample).

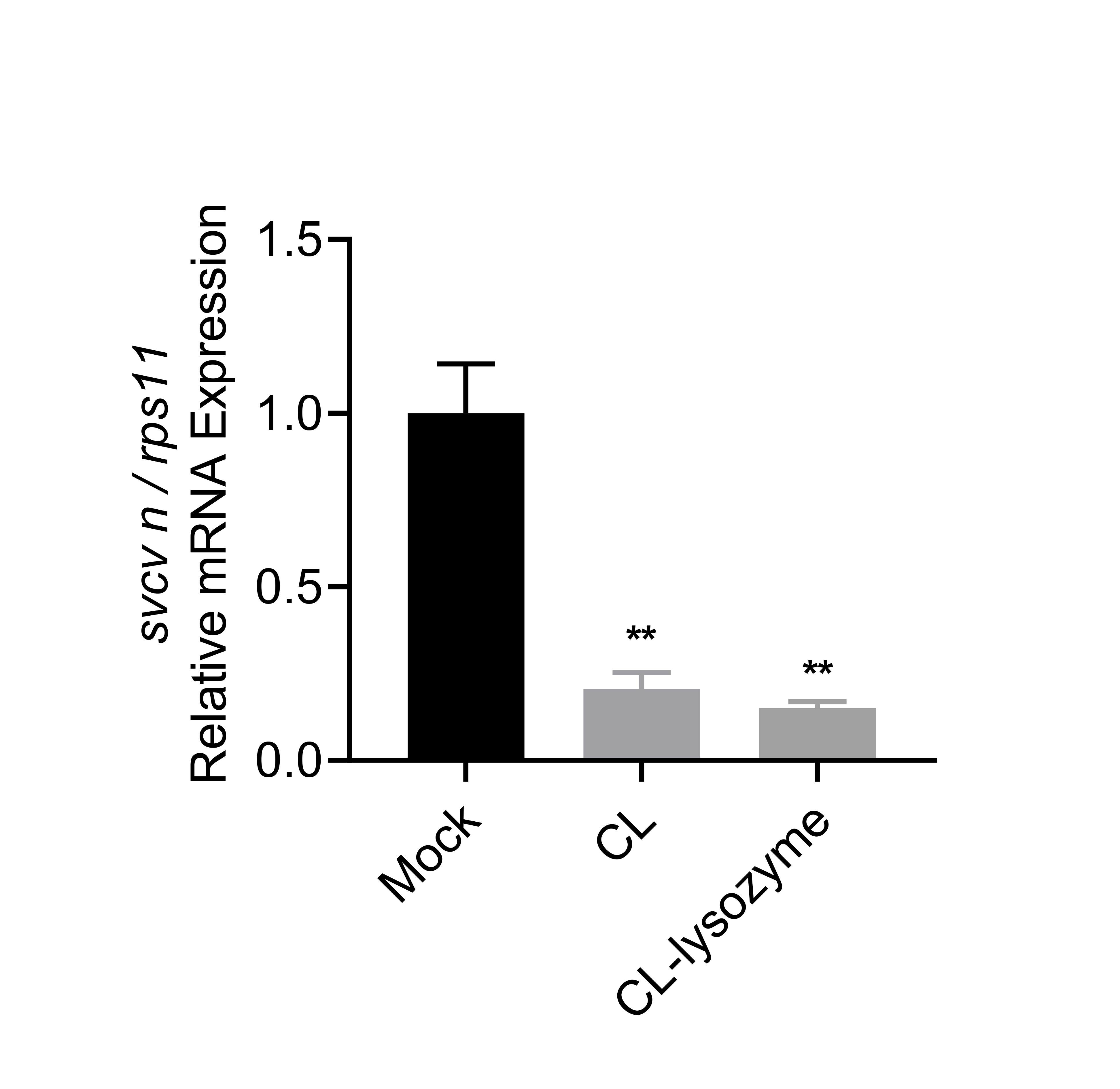

**Fig. S10. Peptidoglycan is not responsible for the antiviral activity of *C. somerae*.** GF zebrafish were treated with cell lysate of *C. somerae* (CL) or lysozyme-treated CL samples and subjected to SVCV infection. Viral replication was detected at 48 hpi. (n = 4, pool of 30 zebrafish larvae per sample).

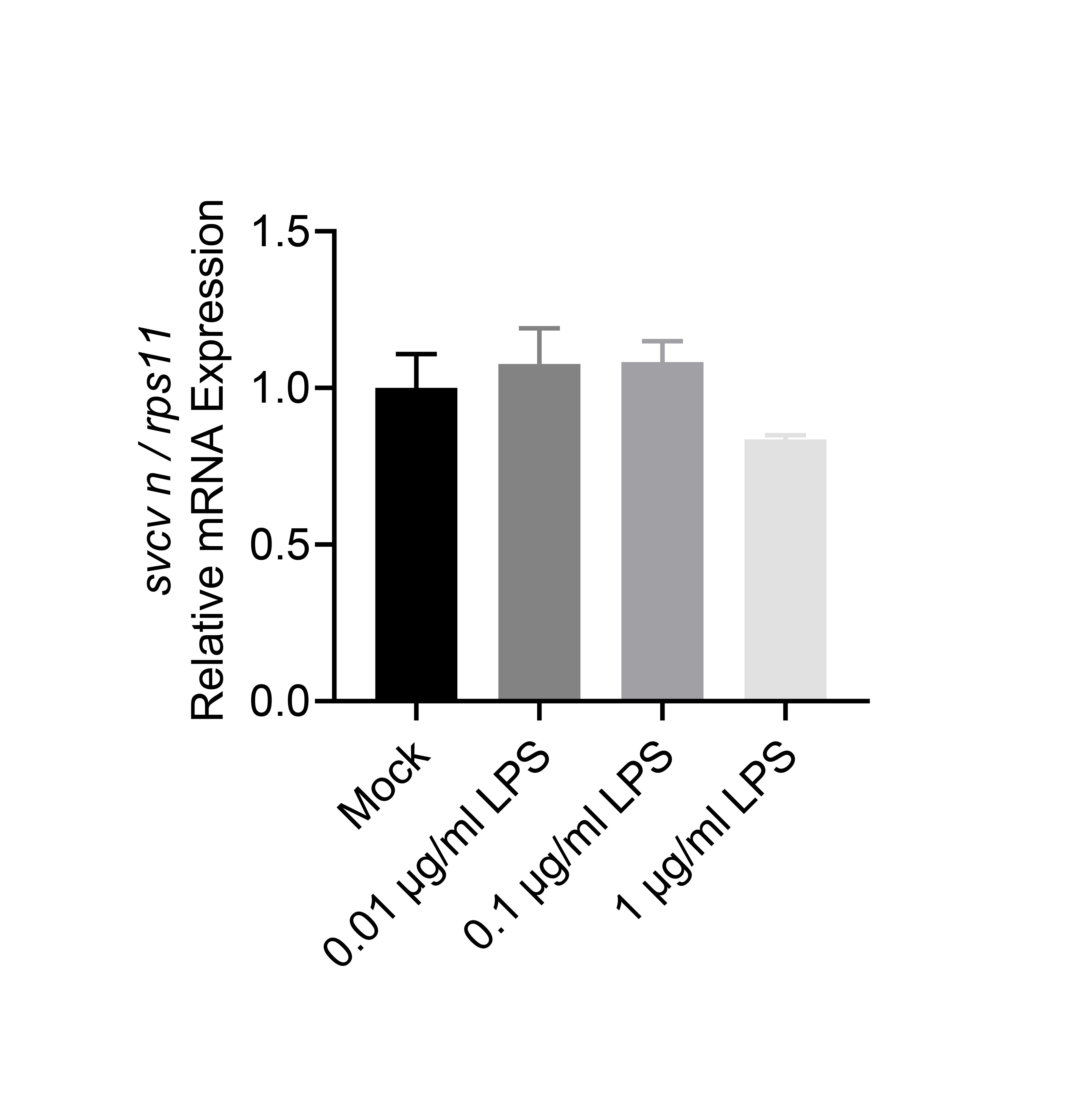

**Fig. S11.** **LPS of *C. somerae* does not have antiviral activity.** LPS was extracted by using a commercial kit (Genmed Scientifics Inc., USA). GF zebrafish were treated with different doses of *C. somerae* LPS and subjected to SVCV infection. Viral replication was detected at 48 hpi. (n = 4, pool of 30 zebrafish larvae per sample).

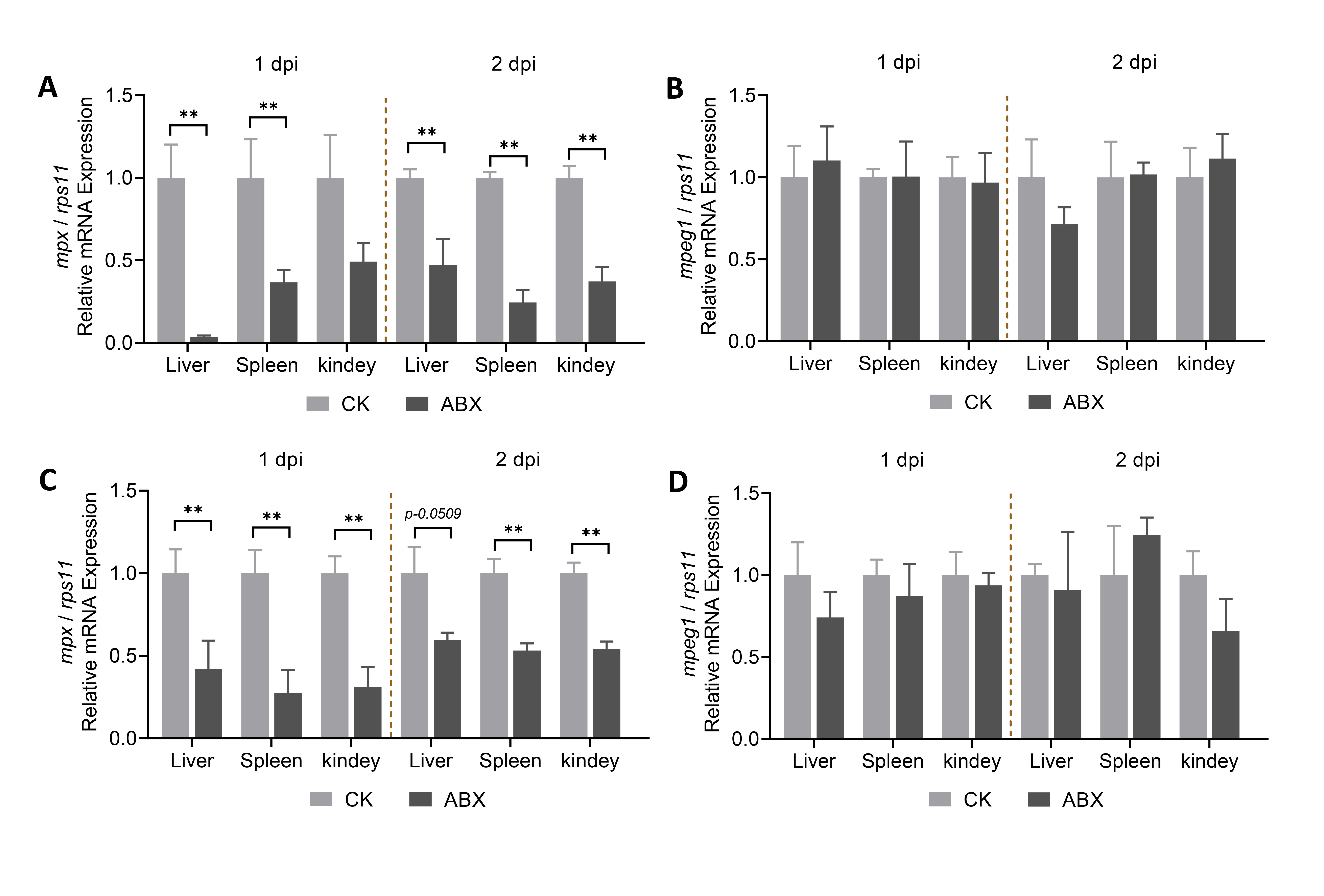

**Fig. S12. Depletion of intestinal microbiome weakens neutrophil response in zebrafish.** (**A**-**B**) Expression of *mpx* (A) and *mpeg1* (B) in the liver, spleen, and kidney of adult zebrafish 1 and 2 days post SVCV infection (n=4, pool of 6 fish larvae per sample). (**C**-**D**) Expression of *mpx* (C) and *mpeg1* (D) in the liver, spleen, and kidney of adult zebrafish 1 and 2 days post poly (I:C) treatment (n=4, pool of 30 zebrafish larvae per sample).
